# Cancer-associated fibroblast compositions change with breast-cancer progression linking S100A4 and PDPN ratios with clinical outcome

**DOI:** 10.1101/2020.01.12.903039

**Authors:** Gil Friedman, Oshrat Levi-Galibov, Eyal David, Chamutal Bornstein, Amir Giladi, Maya Dadiani, Avi Mayo, Coral Halperin, Meirav Pevsner-Fischer, Hagar Lavon, Reinat Nevo, Yaniv Stein, H. Raza Ali, Carlos Caldas, Einav Nili-Gal-Yam, Uri Alon, Ido Amit, Ruth Scherz-Shouval

**Affiliations:** Department of Biomolecular Sciences, The Weizmann Institute of Science, Rehovot, Israel, 76100; Department of Immunology, The Weizmann Institute of Science, Rehovot, Israel, 76100; Chaim Sheba Medical Center, Cancer Research Center, 5262100, Tel-Hashomer, Israel; Department of Molecular Cell Biology, The Weizmann Institute of Science, Rehovot, Israel; Cancer Research UK Cambridge Institute and Department of Oncology, Li Ka Shing Centre, University of Cambridge, Cambridge, UK; Breast Cancer Programme, Cancer Research UK Cancer Centre, NIHR Cambridge Biomedical Research Centre and Cambridge Experimental Cancer Medicine Centre, Cambridge University Hospital NHS Foundation Trust, Cambridge, UK; Chaim Sheba Medical Center, Institute of Oncology, Tel-Hashomer, Israel

**Author notes:** Correspondence should be addressed to I.A. or to R.S.S.

## Abstract

Tumors are supported by cancer-associated fibroblasts (CAFs). CAFs are heterogeneous and carry out distinct cancer-associated functions. Understanding the full repertoire of CAFs and their dynamic changes could improve the precision of cancer treatment. CAFs are usually analyzed at a single time-point using specific markers, and it is therefore unclear whether CAFs display plasticity as tumors evolve. Here, we analyze thousands of CAFs using index and transcriptional single-cell sorting, at several time-points along breast tumor progression in mice, uncovering distinct subpopulations. Strikingly, the transcriptional programs of these subpopulations change over time and in metastases, transitioning from an immune-regulatory program to wound healing and antigen-presentation programs, indicating that CAFs and their functions are dynamic. Two main CAF subpopulations are also found in human breast tumors, where their ratio is associated with disease outcome across subtypes, and is particularly correlated with BRCA mutations in triple-negative breast cancer. These findings indicate that the repertoire of CAFs changes over time in breast cancer progression, with direct clinical implications.

## Introduction

Tumors initiate as a clonal disease, and grow as an ecosystem, in which phenotypically and functionally distinct subpopulations of cells engage in complex interactions that drive tumor progression and metastasis. Genetic and epigenetic heterogeneity among cancer cells flows from the intrinsic biology of multi-step carcinogenesis^1–3^.

Tumors, however, are comprised of more than just cancer cells, and the complexity of tumor heterogeneity is amplified by contributions from the tumor microenvironment^4^. Key players in the tumor microenvironment are cancer-associated fibroblasts (CAFs). CAFs promote cancer phenotypes including proliferation, invasion, extracellular matrix (ECM) remodeling and inflammation^4–7^, as well as chemoresistance^8^ and immunosuppression^9^. Over the years different cell surface markers were shown to identify unique subpopulations of CAFs, and different origins have been suggested for CAFs, including tissue resident fibroblasts, myofibroblasts, bone-marrow (BM) derived mesenchymal stem cells (MSC), and adipocytes^10–14^. The indispensable roles that CAFs play in promoting malignancy and the evident existence of unique subpopulations accentuate the urgency to better understand stromal heterogeneity. Currently, it is unclear to what extent CAFs and their functions change over time with tumor progression and metastasis. These temporal dynamics have been difficult to study, because of two technical hurdles. One is the challenge of sequencing more than a few hundred CAFs, and the second is the lack of cell surface markers for the full range of CAF repertoires, limiting studies to a few known markers, which potentially miss some CAF populations.

Here, we address the question of the full CAF repertoire over time, by using an unbiased approach that does not require a-priori defined markers - massively parallel single cell RNA-sequencing (MARS-seq) and index sorting^15^ - to characterize thousands of CAFs at several time-points over breast tumor growth and metastasis in mice. We identify eight CAF subtypes in two main CAF populations, which we term pCAF and sCAF, based on selective expression of the markers *Pdpn* or *S100a4* (also called fibroblast-specific protein 1; FSP1). These CAF subtypes appear progressively over time, transitioning from an early immune-regulatory transcriptional program, to a late combination of antigen-presentation and wound healing programs. Using the PDPN and S100A4 protein markers, as well as markers for subpopulations of sCAFs and pCAFs, we show that human breast tumors have similar CAF compositions, and that the ratio between the PDPN^+^ and S100A4^+^ CAFs is associated with BRCA mutations in triple-negative breast cancer. Moreover, in two independent cohorts of breast cancer patients, the ratio between the PDPN^+^ and S100A4^+^ CAFs is strongly associated with clinical outcome. This study shows that CAFs functions change with tumor progression, providing clinically relevant markers. Our findings raise the concept of a dynamic tumor microenvironment, in which genomically stable cells change their transcriptional program to keep track of the evolving tumor ecosystem.

## Results

### Comprehensive mapping of breast CAFs reveals subpopulations with distinct transcriptional programs

To discover CAF subtypes that are associated with breast cancer progression we first set out to characterize the stromal cell types/states that comprise breast tumors in a mouse model – triple negative 4T1 cancer cells orthotopically injected into the mammary fat pad of immunocompetent BALB/c mice. This cell line has been extensively used as a robust model for metastatic breast cancer and has been shown to recruit abundant stroma^13^. To expose the full repertoire of stromal transcriptional states and avoid biases driven by a-priori defined cell-type specific markers we used an index sorting and negative-selection based approach for isolation and MARS-seq of CAFs^15^. We densely sampled cells along critical time points of tumor development – at 2 weeks post injection (2W), 4 weeks post injection (4W), and from lung metastases (Met) forming 4-5 weeks post primary tumor injection (and 2-3 weeks after primary tumor resection). Normal mammary fat pad fibroblasts (NMF) from naïve mice were collected as controls. Tumors or normal mammary fat pads were harvested, dissociated into single cell suspensions, and the live cells were stained with the following cell-surface markers: Ter119 (Red blood cells), CD45 (immune cells), and EpCAM (epithelial cells) for negative selection; and Podoplanin (PDPN; Fibroblasts) for index sorting (see Methods). All live cells staining negative for Ter119, CD45 and EpCAM were index sorted and single cell processed by MARS-seq (Fig. 1a; Supplementary Fig. 1a). Overall, we analyzed 8987 QC positive single cells from 12 tumor-bearing mice and 3 control naïve mice (Supplementary Fig. 1b-c; Supplementary Table 1) and used the MetaCell algorithm to identify homogeneous and robust groups of cells (“metacells”; see Methods^16^) resulting in a detailed map of the 88 most transcriptionally distinct subpopulations (Supplementary Table 2). These metacells are organized into 4 broad classes, including: Endothelial cells (characterized by expression of *Pecam1*), pericytes (*Rgs5*), and two classes of fibroblasts that we termed pCAFs (*Pdpn*) and sCAFs (*S100a4*; Supplementary Fig. 1d-e).

**Fig. 1:**
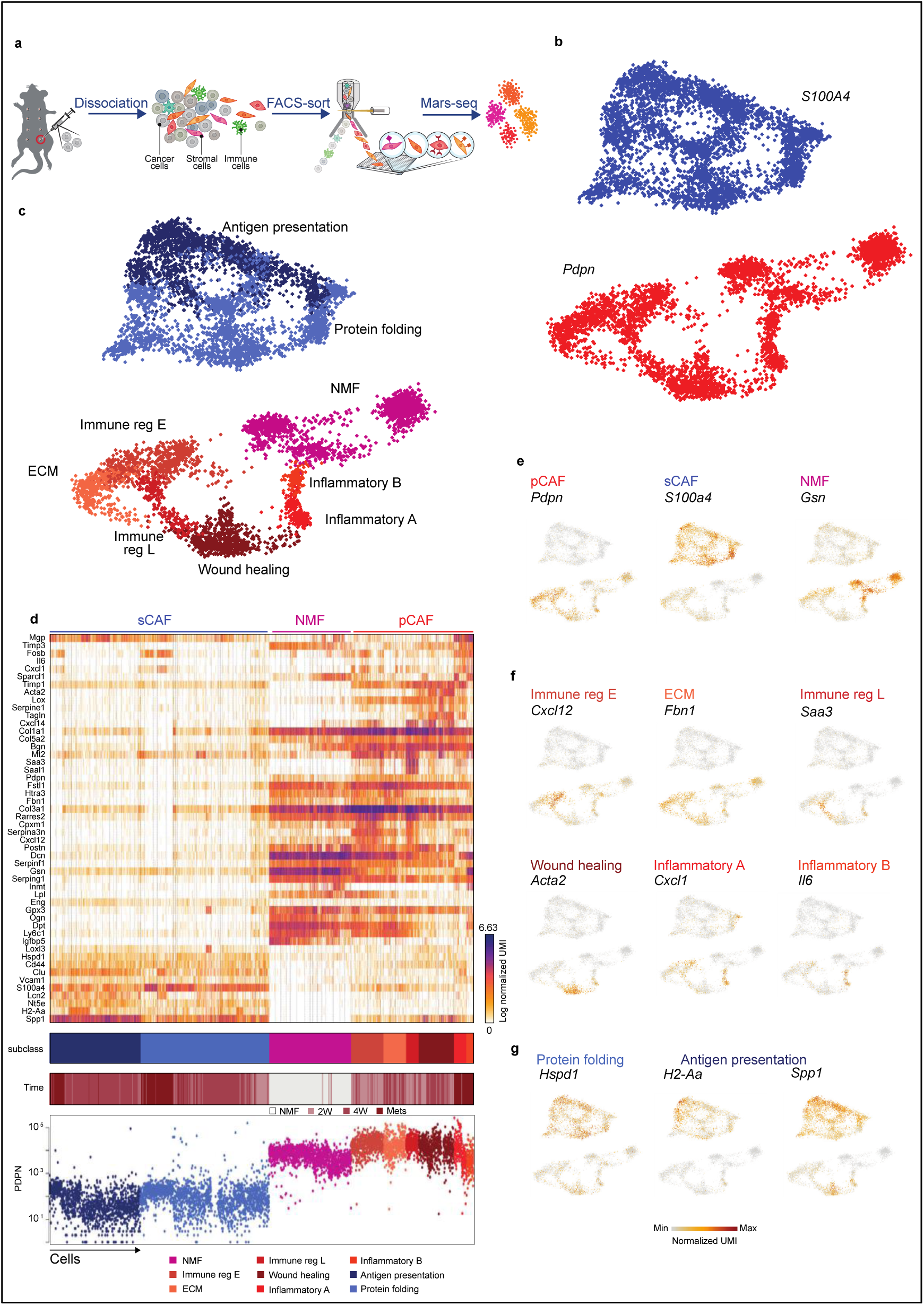
Breast CAFs are comprised of distinct subsets with diverse transcriptional profiles. **a,** Illustration of the experimental procedure. **b and c,** Single cell RNA-seq data from CAF and NMF was analyzed and clustered using the MetaCell algorithm, resulting in a two-dimensional projection of 8033 cells from 15 mice. 83 meta-cells were associated with 2 broad fibroblast populations (**b**) and 9 functional subclasses (**c**) annotated and marked by color code. (**d**) Gene expression of key markers genes across single cells from all subclasses of NMF, pCAF, and sCAF. Lower panels indicate the association to subclass, the time-point, and the PDPN index sorting data, showing protein level intensity in each cell. **e-g,** Expression of key markers genes for NMF, pCAF, and sCAF (**e**); functional annotation for pCAF subclasses (**f**) and sCAF subclasses (**g**) on top of the two-dimensional projection of breast CAFs. Colors indicate log transformed UMI counts normalized to total counts per cell.

We *in silico* removed the *Pecam1* and *Rgs5* cells, as well as a rare population (33 cells) negative for *Pecam1*, *Rgs5, Pdpn,* and *S100a4,* that highly expressed *Myc* and may have originated from cancer cells (see Methods). We continued our analysis with 3875 *Pdpn* cells, and 4158 *S100a4* cells (Fig. 1b; Supplementary Fig. 1d-f; Supplementary Tables 3-4). Each of these broad fibroblast populations could be further divided into functional subsets with distinct transcriptional profiles (Fig. 1c) and differentially expressed genes (Fig. 1d), which were reproducible across mice and batches (Supplementary Fig. 1g). *Pdpn* fibroblasts indeed expressed cell-surface PDPN protein (Fig. 1d lower panel) and included the NMF (*Gsn*) subset (Fig. 1e), and 6 subsets of pCAFs (Fig. 1f). Two of these expressed different gene modules involved in immune regulation and cell migration (*Cxcl12 and Saa3*); one had a wound-healing signature (*Acta2*; encoding for alpha smooth muscle actin; αSMA); one had an extracellular fiber organization signature (*Fbn1*) and two had inflammatory signatures (*Cxcl1 and Il6*). The *S100a4* fibroblasts were completely devoid of NMFs and included 2 subsets of CAFs, albeit these subsets were not as clearly separated from each other as the pCAF subsets (Fig. 1d). One subset, which exhibited high expression of *Spp1* and relatively low expression of *S100a4* (*Spp1^high^S100A4^low^*) was enriched for signatures of antigen presentation (*H2-Aa*) and ECM remodeling (Fig. 1g). The other population exhibited high expression of *S100a4* and relatively low expression of *Spp1* (*Spp1^low^S100A4^high^*) and was enriched in protein folding and metabolic genes (*Hspd1*; Fig. 1g). To validate our single cell sequencing results we performed bulk RNA sequencing of sCAFs, pCAFs, and NMFs, and compared the profiles obtained by bulk and single-cell RNA sequencing. All groups showed high correlation (R>0.5) between bulk and cognate single cell profiles (Supplementary Fig. 1h). We also found high correlation between pCAF and NMF profiles. sCAFs, however, showed no correlation with either pCAFs or NMFs (Supplementary Fig. 1h), suggesting that they have further diverged from NMF (as compared to pCAFs), or perhaps have a different origin altogether. Cancer cells that have undergone EMT were recently shown to contribute to the stromal milieu in mice^14^. In order to exclude their potential contribution to the sCAF population, we analyzed the bulk RNA sequencing data for lineage traces of 4T1 cancer cells (transfected plasmid reads; see Methods section for details). This analysis confirmed that while some cancer cells may have escaped the negative selection approach, the majority of sCAFs are derived from host mesenchymal cells (Supplementary Table 5; see Methods for details).

### CAF composition is dynamically reshaped as tumors progress and metastasize

Tumor heterogeneity increases with tumor progression^1, 17, 18^. Similarly, we and others have hypothesized that stromal heterogeneity increases as tumors progress. Accordingly, our analysis shows that metacell composition varies extensively between the different time points (Fig. 1d and Fig. 2a-b). Normal mammary fat pads harbored *Pdpn^+^* fibroblasts and were devoid of *S100a4^+^* fibroblasts. Two weeks after tumor initiation (2W) significant heterogeneity is observed: sCAFs, which were absent at the time of injection, now constitute ∼30% of the CAF population (Fig. 2b). The majority of sCAF express metabolic and protein folding genes (*Hspd1*). The remaining ∼70% of CAFs at 2W are *Pdpn^+^*, yet in stark contrast to *Pdpn^+^* NMF, pCAFs are highly heterogeneous – more than half of them belong to the two immune regulatory subpopulations (*Cxcl12* or *Saa3*), ∼10% of them express ECM organizing modules (*Fbn1*), and the remaining quarter exhibits a wound healing transcriptional profile (*Acta2*). 4 weeks after tumor initiation (4W) the majority of CAFs are sCAFs (∼77%) while only ∼23% are pCAFs. Once again, the composition of metacells within each class of CAFs has changed dramatically - The dominant pCAF populations in 4W tumors are the wound healing class (*Acta2*) and ECM organizing pCAFs (*Fbn1*), while the immune regulatory pCAF subpopulations (*Cxcl12; Saa3*) are diminished (Fig. 2b). sCAF at 4W are composed largely of cells expressing ECM remodeling and antigen present (*Spp1, H2-Aa*). Lung metastases (Mets) spontaneously arising from these tumors contain mostly sCAFs (∼70%) and share similar sCAF subpopulations with primary tumors (i.e. *Spp1 and Ha-Aa; Hspd1*), at 1:1 ratios. The pCAF population in Mets (∼30%) is comprised mostly of two unique inflammatory subpopulations of pCAF (*Il6* and *Cxcl1*) that were not observed in the primary tumors or in the normal mammary fat pad (Fig. 2b). The dynamic shift in CAF composition was confirmed by FACS analysis of cell-surface PDPN protein expression in the different time points. As tumors grew, the abundance of PDPN*^+^* cells within the stromal (CD45^-^EpCAM^-^) population decreased, and the abundance of PDPN^-^ cells increased (Supplementary Fig. 2a).

**Fig. 2:**
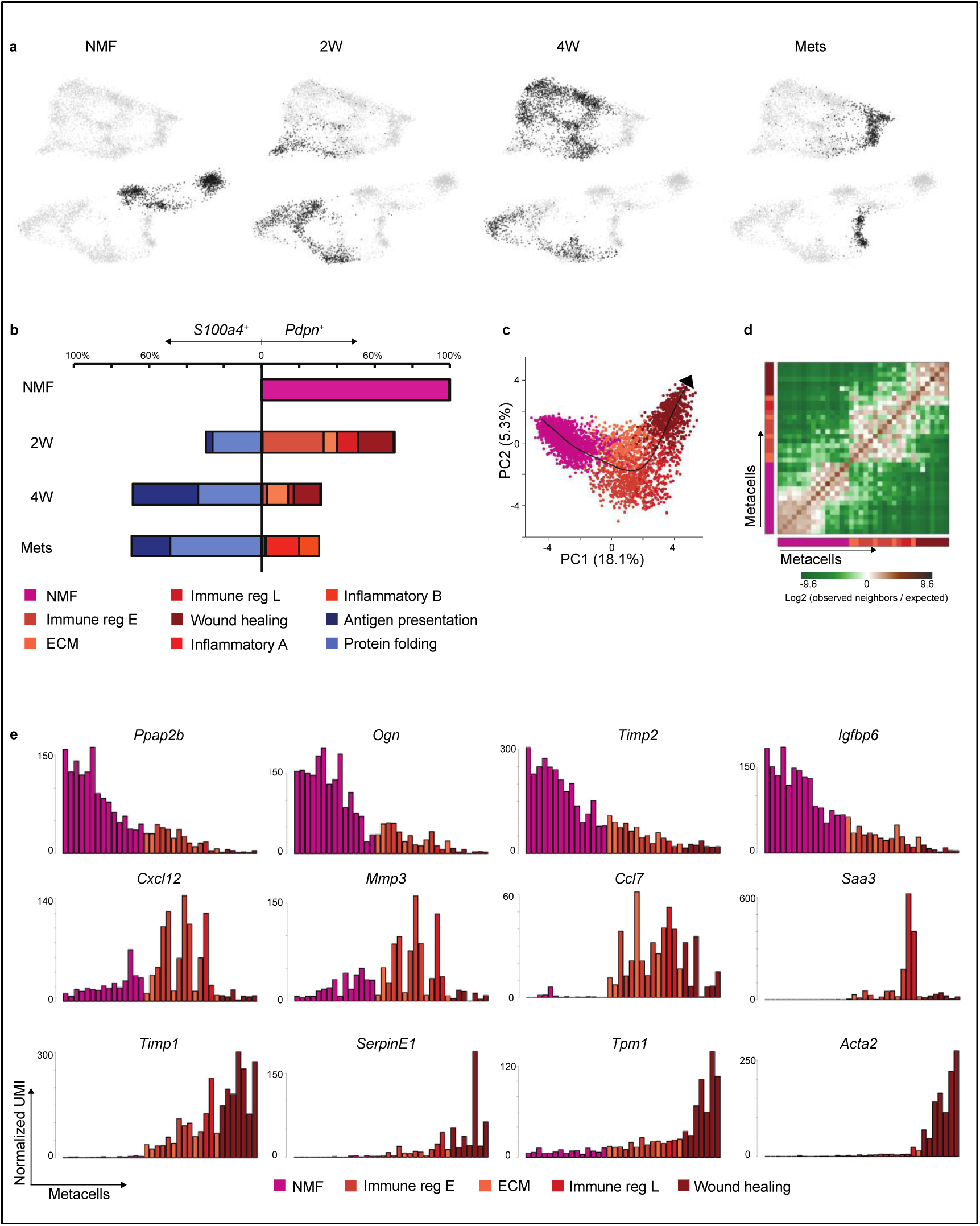
CAF composition and gene expression changes with tumor growth and metastasis. **a,** Projection of cells from different time points (black) on top of the 2D map of breast fibroblasts (presented in Fig. 1b-c). **b,** Compositions of *Pdpn^+^* fibroblasts (right) and *S100a4^+^* fibroblasts (left) at different time points (normalized to 100% total fibroblasts). Subclasses are annotated and color-coded. **c-e,** Slingshot analysis of pseudo-time trajectory from NMF to pCAF from 2W and 4W. Cells are color-coded as in **b**. **c,** Suggested trajectory from NMF to pCAF projected over the top two principal components. **d,** Heat map showing enrichment (log2 fold change) for kNN connections between metacells over their expected distribution. Metacells are ordered by their position on the Slingshot pseudotime. **e,** Expression of hallmark NMF and pCAF genes across metacells (average UMI/cell), ordered by pseudo-time.

### *Pdpn^+^* fibroblasts diverge into pro-tumorigenic CAFs during tumor progression

Different origins have been proposed for CAFs, including normal tissue fibroblasts, mesenchymal stromal cells, and adipocytes^10–13^. Our metacell analysis showed that pCAFs (but not sCAFs) share similar patterns of transcription with NMFs (Fig. 1d and Supplementary Fig. 1h), suggesting that pCAFs may have originated from NMFs. To infer the most probable transcriptional trajectory for pCAFs we applied Slingshot, a computational method for cell lineage pseudo-time inference^19^. Slingshot analysis displayed a gradual transition from NMFs through early immune regulatory and ECM organizing pCAFs, to late immune regulatory pCAFs, and eventually to wound healing pCAFs (Fig. 2c-d). This trajectory is consistent with the transition from normal fibroblasts through 2W tumors to 4W tumors (Supplementary Fig. 2b-c). NMFs expressed high levels of hallmark genes encoding membrane bound and extracellular proteins (*Ppap2b*, *Ogn*, *Timp2, Igfbp6*, *Igfbp5*, *Dpp4*). Expression of these signature genes gradually decreased along the trajectory leading to wound-healing pCAF (Fig. 2e, upper row, and Supplementary Fig. 2d). In parallel, gradual increase in expression was observed along this trajectory for signature genes involved in cell migration and wound healing, such as *Timp1, Serpine1, Tpm1 and Acta2* (Fig. 2e, lower row, and Supplementary Fig. 2d). A third pattern observed along this trajectory was that of genes whose expression was low in NMFs, high in the ECM/immune-regulatory pCAFs and low again in wound-healing pCAFs. These included genes such as *Cxcl12*, *Mmp3, Ccl7, and Saa3* (Fig. 2e, middle row, and Supplementary Fig. 2d).

### sCAFs are transcriptionally distinct from pCAFs and NMFs

sCAFs exhibit global gene expression profiles that are different from those of pCAFs and NMFs (Supplementary Fig. 1f). Moreover, we could not find transitional cells linking these fibroblast types (Fig. 1d) that would suggest a gradual shift from NMFs to sCAFs, as we observe for pCAFs. Bone marrow (BM) derived mesenchymal stromal cells (MSCs) are commonly viewed as a source of CAFs^10, 20^. The molecular chaperone Clusterin was recently shown to play a tumor-promoting role in BM-MSC derived CAFs recruited to breast tumors in mice^10^. Indeed, Clusterin (*Clu*), as well as several other MSC markers (*Vcam1, Cd44*, *Eng*, and *Nt5e*), were differentially upregulated in sCAFs compared to pCAFs (Fig. 1d and Fig. 3a). Taken together with the observation that *S100a4*^+^ fibroblasts are not found in the normal mammary fat pad, this may suggest that sCAFs arise from a different origin than pCAFs, and are recruited to the tumor, perhaps from BM-MSCs.

**Fig. 3:**
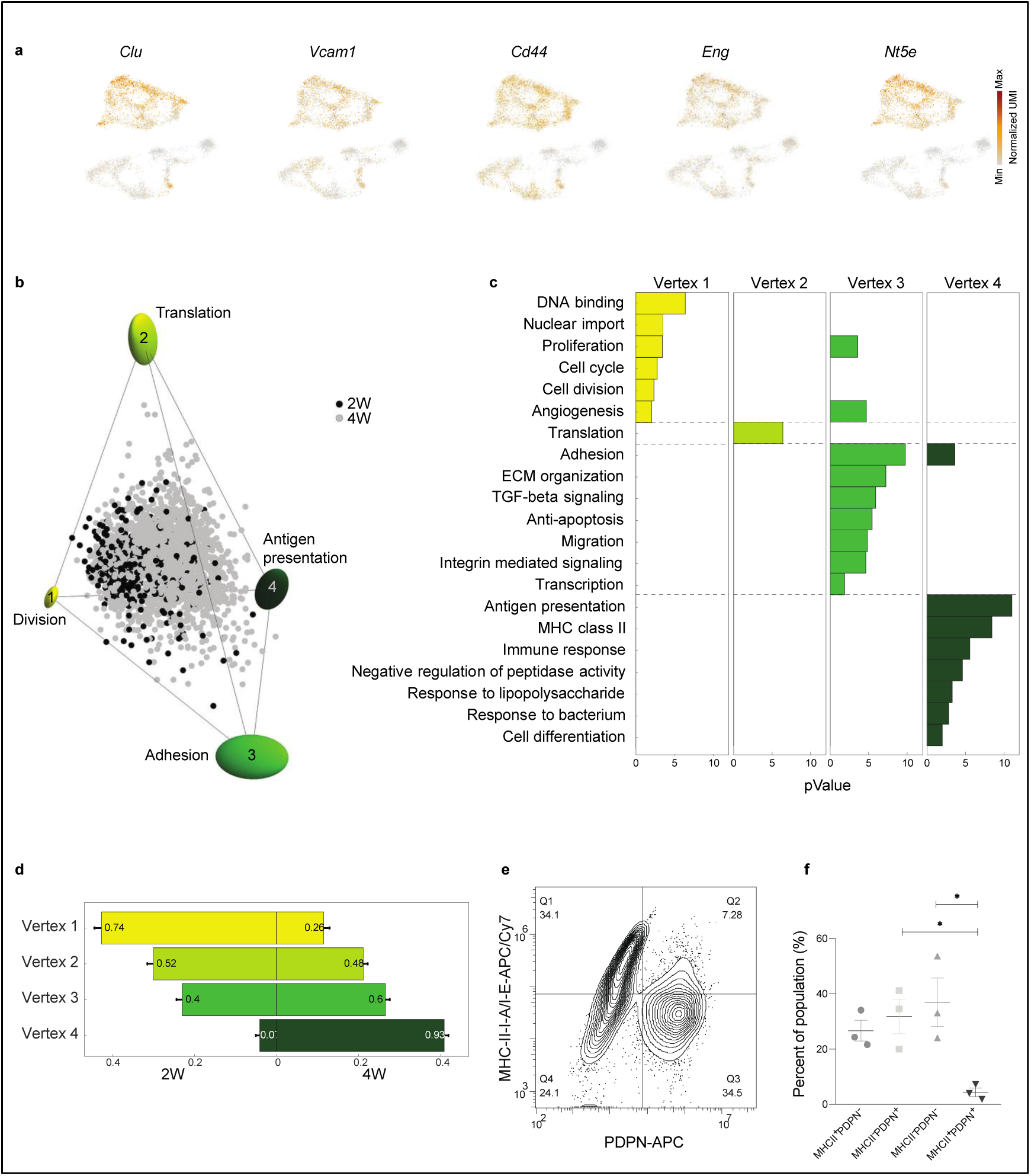
sCAFs show a continuum of cell states which fills a tetrahedron in gene-expression space, suggesting trade-off between 4 functions. **a,** Expression of hallmark mesenchymal stem cell marker genes on top of the two-dimensional projection of breast cancer stroma (presented in Fig. 1b-c). Colors indicate log transformed UMI counts normalized to total counts per cell. **b,** ParTI analysis of 2W and 4W sCAF single-cell gene-expression in the space of the first 3 principal components shows a continuum that can be well enclosed by a tetrahedron. At the vertices are ellipses that indicate standard deviation of vertex position from bootstrapping. Cells are color-coded according to time point. Vertices are annotated and color-coded. **c,** Gene ontology enrichment in the different vertices (see full list in Supplementary Table 6). **d,** Relative representation of each time point in the 4 vertices. The x-axis shows the fraction of cells from 2W and 4W closest to each vertex. Numbers in the bars are the fraction of each time point in the 100 cells closest to each archetype. **e-f,** Flow cytometry analysis of cell surface expression of MHC-II molecules I-A/I-E vs PDPN in CD45^-^EpCAM^-^ cells from 4W tumors. A representative flow cytometry plot is shown in (**e**), quantification of results from 3 biological replicates is presented in (**f**) as average ± SEM. P-values were calculated using one-way Anova correcting for multiple comparisons. * p<0.05

### sCAFs show a continuum of cell states bounded by four major transcriptional programs

Unlike pCAFs, the sCAFs do not seem to form discrete subpopulations, but rather a continuum in gene expression space, which implies a continuum of cell states. To infer biological functions associated with these cell states, we applied the Pareto task inference (ParTI) method^21, 22^. The method is based on an evolutionary theory suggesting that when cells need to perform multiple functions, no single gene-expression profile can be optimal for all functions at once. This trade-off leads to specific patterns in the data: individual cells fall into a polyhedron in gene expression space^22^. At the vertices of the polyhedron are the gene expression profiles optimal for each of the functions. Cells near a vertex are specialists at that function, whereas cells near the middle of the polyhedron are generalists^22, 23^. We first applied ParTI on NMFs, 2W, and 4W CAFs. Mets clustered separately in this analysis and were therefore excluded (see Methods). pCAFs clustered with NMFs, and sCAFs formed a distinct cluster, confirming our metacell analysis (Supplementary Fig. 3a). Next we analyzed each cluster separately. pCAFs and NMFs formed a 1D continuum (a curve) in agreement with the slingshot analysis (Supplementary Fig. 3b). sCAF gene expression could not be explained well by a continuum in 1D (curve) or 2D (a planar polygon; Supplementary Fig. 3c). Rather, their transcriptional states were best described as a continuum in a tetrahedron (Fig. 3b, Supplementary Fig. 3d-i). At the vertices of this tetrahedron are 4 transcriptional programs representing distinct biological functions (Fig. 3b and Supplementary Fig. 3d): Vertex 1 is enriched with cells that express programs for cell division and proliferation (Fig. 3c and Supplementary Table 6). Vertex 2 corresponds to protein translation. Cells near vertex 3 express adhesion, ECM organization, pro-survival and migration programs. Vertex 4 corresponds to immune response programs, in particular antigen presentation via MHC class II genes (Fig. 3c and Supplementary Table 6). The distribution of cells within the tetrahedron changed with tumor growth (Fig. 3d). sCAFs in 2W tumors were located mostly in the space between vertices 1-3, indicating that they express transcriptional programs of division, adhesion, and protein translation (Fig. 3d). Antigen presentation programs were expressed mostly by 4W sCAFs, and scarcely by 2W sCAFs, suggesting a temporally dynamic division of functions between sCAFs in breast tumors.

To further test the finding that a subset of sCAFs expresses MHC class II proteins we performed flow cytometry analysis of sCAFs and pCAFs from 4W tumors, and compared the percentage of I-A/I-E^+^ cells in the two CAF subsets. Whereas pCAFs scarcely expressed MHC-II, these MHC class II cell-surface molecules were expressed by ∼50% of 4W sCAFs (Fig. 3e-f).

### PDPN and S100A4 mark mutually exclusive, morphologically distinct CAFs in mouse breast tumors

To validate our classification and examine the spatial distribution of the CAF subpopulations that we have identified, we performed immunohistochemical staining of 4T1 tumors from different stages with anti-S100A4 and anti-PDPN antibodies. Cytokeratin (CK) was used as a marker for cancer cells. The normal mammary fat pad showed very weak expression of S100A4, whereas PDPN^+^ fibroblasts were highly abundant (Fig. 4a, upper panel). 2W and 4W tumors harbored both PDPN and S100A4-positive cells, and the expression pattern of both proteins was different than that of CK, suggesting that these are stromal cells (Fig. 4a, middle panels). Metastases were rich in S100A4-positive cells (Fig. 4a, lower panel). PDPN, however, was scarcely expressed in the metastatic region, and strongly expressed in the normal adjacent lung tissue (Fig. 4a, lower panel). At all tumor stages that we monitored, pCAFs were long and spindly, resembling the morphology of NMFs, and sCAFs were smaller. Both classes of CAFs were distributed in all regions of the tumor, and we could not identify a distinct spatial pattern of distribution within the tumor (Fig. 4a).

**Fig. 4:**
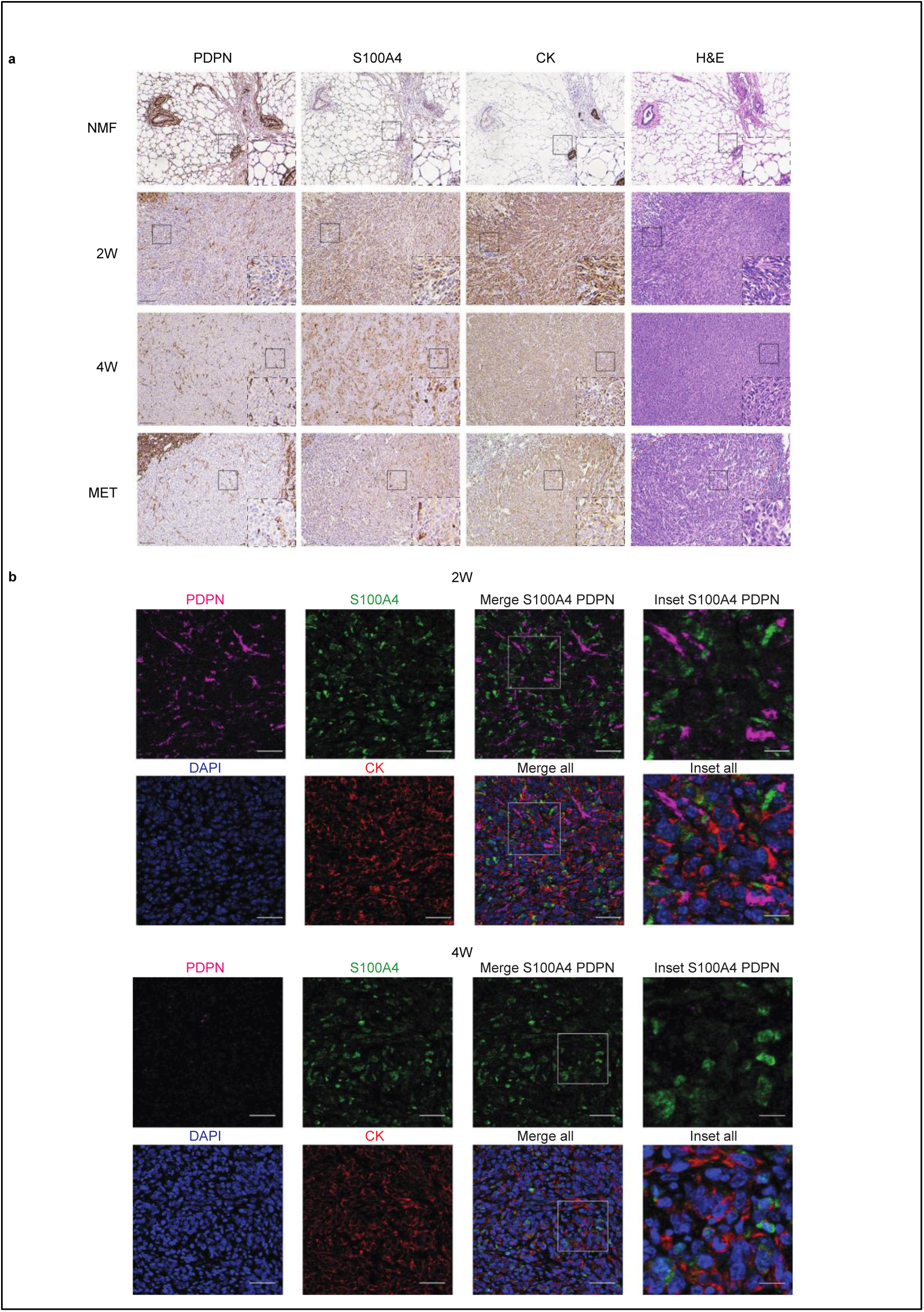
PDPN and S100A4 proteins are expressed on distinct types of breast CAFs in mouse tumors. **a,** Consecutive formalin fixed paraffin embedded (FFPE) tissue sections of tumors, metastases, or normal mammary fat pads were immunostained with antibodies against the indicated proteins, or stained with hematoxylin & eosin (H&E). Staining was performed on 3 independent biological replicates for each time point; Representative images are shown. All images were collected at the same magnification and are presented at the same size. Scale bar = 100μm. For each panel, regions marked by rectangles are shown as 2.5X insets in black dashed rectangles. A dashed red line on the H&E marks the metastatic region in the lung. **b,** Multiplexed immunofluorescent (MxIF) staining was performed on all biological replicates from (**a**) with antibodies against the indicated proteins. Representative images of 2W and 4W tumor FFPE sections are shown. Scale bar = 50 μm, inset scale bar = 17μm.

Multiplexed immunofluorescent (MxIF) staining confirmed that S100A4 and PDPN mark different populations of cells (Fig. 4b and Supplementary Fig. 4). We saw partial overlap between S100A4 and CK staining, mostly in normal mammary fat pads (Supplementary Fig. 4a and c). These results suggest that mostly normal epithelial cells but to a low degree also cancer cells may express S100A4 (Supplementary Fig. 4c). Nevertheless, the majority of S100A4 cells in primary tumors and in metastases were CK-negative, confirming our sequencing results and suggesting that PDPN^+^ cells and S100A4^+^ cells are distinct subtypes of CAFs (Fig. 4b and Supplementary Fig. 4).

### Ly6C^+^ pCAFs are immunosuppressive

Our sequencing results suggested that pCAFs are comprised of diverse subpopulations performing distinct tasks such as immune regulation and wound-healing. To test the functional relevance of these findings we isolated pCAFs from 4T1 mouse tumors using FACS sorting with PDPN as a positive selection marker, and then further separated these cells into subpopulations using Ly6C as a marker for the immune regulatory subpopulation and SMA (encoded by *Acta2*) as a marker for the wound-healing subpopulation (Fig. 1d). The two proteins were mutually exclusive and marked distinct subpopulations of cells (Fig. 5a). The Ly6C^+^SMA^-^ subpopulation was most abundant in the NMFs and decreased as tumors progressed, while the Ly6C^-^SMA^+^ subpopulation showed the opposite trend – it was lowest in NMFs and increased as tumors progressed (Fig. 5b). This trend was shared also with a Ly6C^-^SMA^-^ subpopulation.

**Fig. 5:**
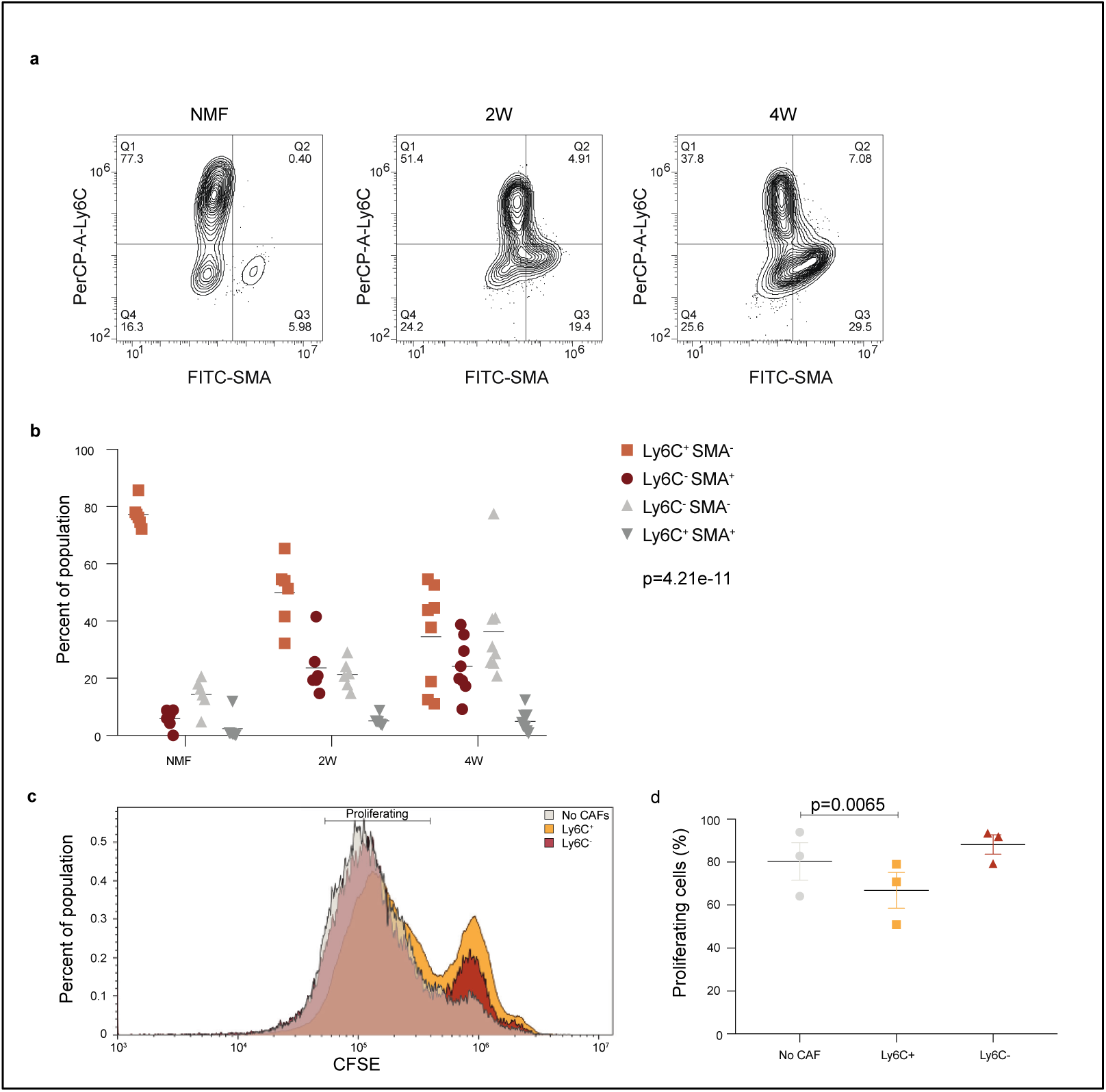
Ly6C^+^ pCAFs suppress CD8 T-cell proliferation, *in vitro*. a-b, FACS analysis of Ly6C and SMA expression in CD45^-^EpCAM^-^PDPN^+^ cells freshly harvested from normal mammary fat pads, 2W tumors, and 4W tumors, and immediately fixed. Representative flow cytometry plots are shown in (**a**) and the results from 3 independent experiments are quantified and analyzed utilizing two-wayAnova in (**b**). **c-d,** CD45^-^EpCAM^-^PDPN^+^ cells from 4W tumors were sorted to Ly61^+^ vs Ly6C^-^ populations, which were then incubated in vitro at 1:1 ratios with CD8 T cells activated by CD3/CD28 beads and marked by CFSE. Representative FACS plots of CFSE signals after 36h of co-incubation are shown in (**c**) and the results from 3 independent experiments are quantified and analyzed utilizing Anova in (**d**).

Since Ly6C^+^SMA^-^ pCAFs expressed a module of immune regulatory genes we examined their ability to suppress T cell activation, *in vitro*. While Ly6C^-^SMA^+^ pCAFs had no significant effect on T cell proliferation, Ly6C^+^SMA^-^ pCAFs significantly suppressed CD3/CD28-mediated T cell proliferation (Fig. 5c-d). These results support our molecular profiling results and suggest that Ly6C^+^SMA^-^ pCAFs are immune regulatory, while Ly6C^-^SMA^+^ are not.

### S100A4 and PDPN mark distinct stromal populations in human breast tumors

To test the clinical relevance of our findings we performed MxIF staining for PDPN and S100A4 in human estrogen receptor positive (ER^+^) and triple negative (TN) breast cancer patient tissue samples. CK staining was performed to mark epithelial cancer cells (Fig. 6a). We found that PDPN^+^ cells and S100A4^+^ cells are major constituents of human breast cancer stroma, and exhibited very low overlap with CK staining (Fig. 6a and Supplementary Fig. 5f-g). A minor overlap was observed between S100A4 and CD45 staining, in cells with mesenchymal morphology (Supplementary Fig. 5a). Similar to our mouse model, PDPN^+^ cells and S100A4^+^ cells were mutually exclusive. These observations suggest that PDPN and S100A4 mark distinct subtypes of CAFs in human breast tumors. We observed partial segregation in the spatial organization of CAFs in human tumors (Fig. 6a). Both in ER^+^ and TN samples, a subset of pCAF was found immediately adjacent to CK^+^ cancer cells, or infiltrating the cancerous region. The infiltrating pCAFs were less spindly than the elongated pCAFs found in stromal stretches. The rest of the pCAFs were dispersed in stromal regions, mixed with sCAFs. In contrast, sCAFs were less frequently found immediately adjacent to cancer cells (Fig. 6a, insets).

**Fig. 6:**
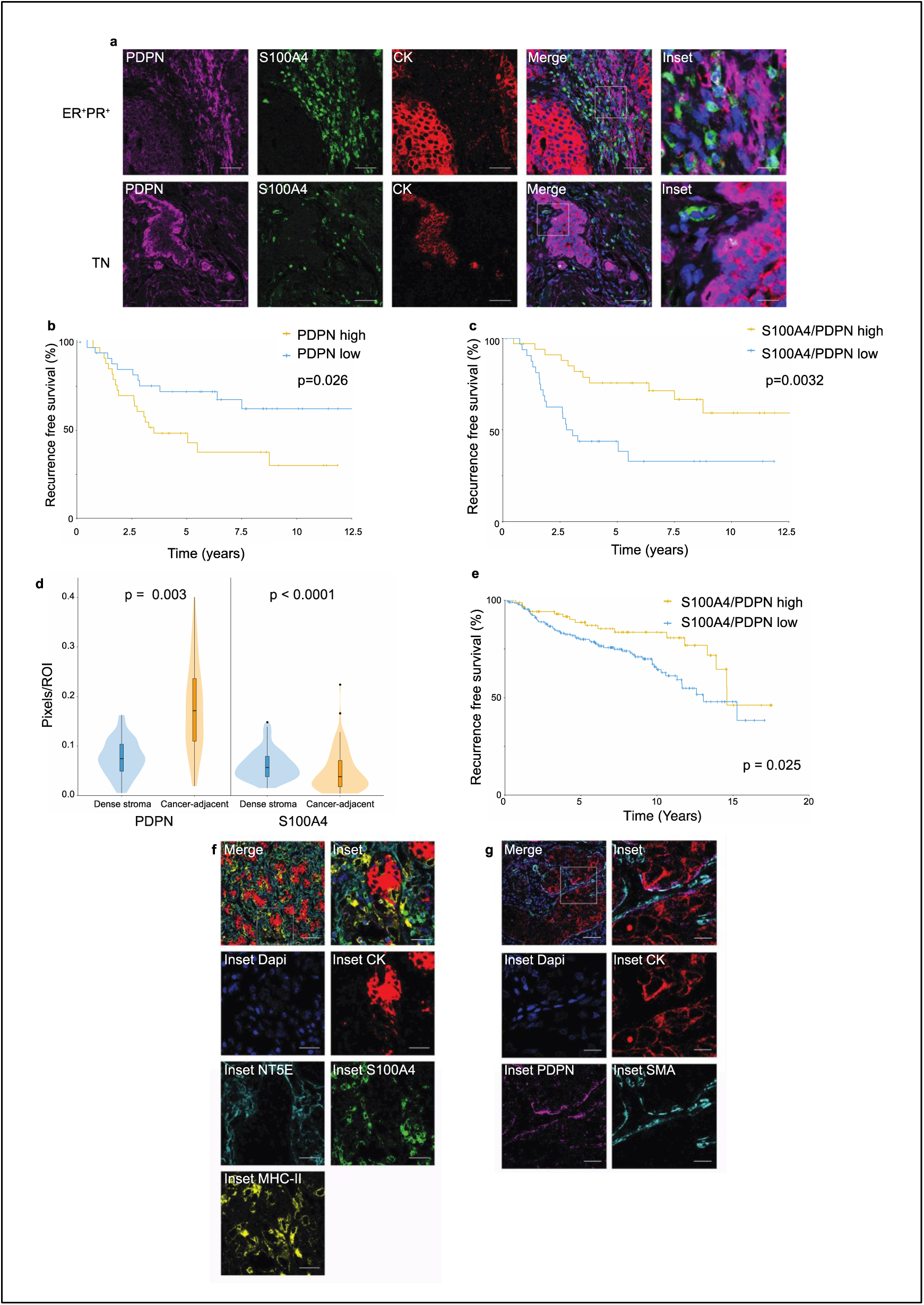
PDPN and S100A4 stromal staining is correlated with disease outcome in human breast cancer patients. **a,** MxIF staining of FFPE tissue sections from ER^+^ or TN breast cancer (BC) patients with antibodies against the indicated proteins. Staining was performed on 5 ER^+^ and 6 TN patients, representative images from an ER^+^PR^+^HER2^-^ and a TN patient are shown (each row is a different patient). Scale bar = 50 μm; inset scale bar = 12.5 μm. **b-c,** FFPE tumor microarray (TMA) sections from a cohort of 72 TNBC patients were immunostained for PDPN and S100A4 and scored (see Methods section). PDPN scores (**b**) or S100A4/PDPN scores (**c**) were classified as higher or lower than the median, and the association with recurrence-free survival was assessed by Kaplan Meier (KM) analysis. P-value was calculated using log rank test. **d,** Cancer-adjacent regions and regions of dense stroma were determined for each core in the TNBC TMA based on CK staining (see Methods section), and PDPN and S100A4 staining in each region was scored. P-value was calculated using Wilcoxon matched pairs signed rank test. **e,** FFPE TMA sections of 293 breast cancer patients from the METABRIC cohort were stained and scored for PDPN and S100A4 as described in (**b-c**). S100A4/PDPN scores were classified as higher (n=88) or lower (n=200) than 1 (5 outlier samples were omitted from the analysis; see Methods section), and the association with recurrence-free survival was assessed by Kaplan Meier (KM) analysis. **f-g,** Full tumor FFPE sections from a subset of 12 patients of the TNBC cohort were MxIF stained with antibodies against the indicated proteins. Representative merged images are shown, with merged insets and insets of the independent channels (**f and g** are each of a different patient). Scale bar = 50 μm; inset scale bar = 20 μm.

### S100A4/PDPN ratio is correlated with disease outcome in two independent cohorts of breast cancer patients

To study the clinical significance of these findings, we co-stained and scored PDPN, S100A4, and CK immunostaining in a cohort of 72 TN breast cancer (TNBC) patients with documented long-term clinical follow-up (Supplementary Table 7). For each patient, we stained 3 cores of the tumor, calculated the average area of positive staining for each marker (Supplementary Fig. 5b), as well as ratios between the 3 markers, and evaluated whether these staining scores correlate with each other (Supplementary Fig. 5c), and with disease outcome. High CK expression levels led to increased hazard of recurrence, as expected, and significantly correlated with poor survival (p=0.028; Supplementary Table 8). Next we evaluated our stromal markers. We found that PDPN levels significantly correlated with disease outcome (p=0.013, Supplementary Table 8): patients whose tumors had high PDPN levels had shorter recurrence-free survival (p=0.026, Fig. 6b), as well as overall survival (p=0.0011, Supplementary Fig. 5d). S100A4 on its own was not significantly correlated with disease outcome in this cohort of patients, yet it showed an opposite hazard ratio to that of PDPN (Supplementary Table 8). We therefore asked whether evaluation of the S100A4/PDPN ratio could improve our ability to predict patient outcome. Indeed, we observed a striking correlation between high S100A4/PDPN ratios and increased recurrence-free survival (p=0.0032) and overall survival (p=0.00015; Fig. 6c and Supplementary Fig. 5e).

To quantitatively test the observation that pCAFs infiltrate the cancerous region more often than sCAFs we defined regions of dense stroma versus cancer-adjacent regions based on CK staining, and calculated the average area of positive staining for S100A4 and PDPN in each region (Supplementary Fig. 6a). pCAFs were significantly more abundant in cancer-adjacent (ca) regions than in dense stroma (ds) regions (Fig. 6d). The average ratio of caPDPN/dsPDPN staining was 3.0 (Supplementary Fig. 6b). sCAFs exhibited a starkly different distribution. They infiltrated the cancerous region significantly less than pCAFs, and the average ratio of caS100A4/dsS100A4 was 0.8 (Fig. 6d and Supplementary Fig. 6b).

Our initial observation that S100A4 and PDPN stain not only TNBC but also ER^+^ breast cancer patient samples suggested that S100A4/PDPN ratio may be a general marker of disease outcome in breast cancer. To test this hypothesis we repeated our staining and scoring of PDPN, S100A4, and CK in an independent cohort of 293 breast cancer patients with documented long-term clinical follow-up and molecular profiling data from the METABRIC study^24^ (Supplementary Table 9). In this cohort of mixed breast cancer subtypes, S100A4/PDPN ratios significantly correlated with disease progression (p=0.025, Fig. 6e). Similar to our results from the TNBC cohort, high S100A4/PDPN ratios were associated with increased recurrence free survival in the METABRIC cohort (Fig. 6e). The spatial distribution of sCAFs and pCAFs was also similar in the METABRIC cohort to that of the TNBC cohort. Namely, the average ratio of caPDPN/dsPDPN staining was significantly higher than the average ratio of caS100A4/dsS100A4 (Supplementary Fig. 6c-d).

### A subset of human sCAFs expresses MHC class II, whereas a subset of pCAFs expresses SMA

To further characterize human sCAFs and pCAFs, we tested several of the markers for sCAF and pCAF subpopulations found in our scRNA-seq data by MxIF in a subset of the TNBC cohort. NT5E (aka CD73) and the MHC class II cell-surface receptor HLA-DR each marked a distinct subset of sCAFs, with very low overlap (Fig. 6f and Supplementary Fig. 5f). SMA was expressed by a subset of pCAFs (Fig. 6g and Supplementary Fig. 5g). These results support our findings from the 4T1 murine model and provide new combinations of markers to detect distinct CAF subpopulations in human patients. To test for possible correlation between S100A4/PDPN ratio and T-cell infiltration we next stained and scored CD3 in our TNBC cohort (Supplementary Fig. 7a-b). We found no significant correlation between CD3 and disease outcome (Supplementary Table 8), nor did CD3 staining correlate with any of the other cell markers tested (CK, PDPN, S100A4; Supplementary Fig. 5c).

### High S100A4/PDPN ratios are associated with BRCA mutations in TNBC

A substantial fraction of TNBC patients carry mutations in BRCA genes (in particular BRCA1^25^), and BRCA mutations frequently lead to TNBC^26^. These mutations are particularly prevalent in the Israeli population (specifically in Ashkenazi Jews). While the METABRIC cohort had very few BRCA mutated patients, in our TNBC cohort 20/45 patients (for which BRCA status was documented) carried such mutations (Supplementary Tables 7, 9). These patients exhibited increased T-cell infiltration (as measured by CD3 staining) compared to patients with WT BRCA (Supplementary Fig. 7c), yet neither T-cell infiltration nor BRCA status correlated with survival (Supplementary Fig. 7d and Supplementary Table 8 and 10).

We therefore tested for possible associations between BRCA status, CAF marker expression, and survival (Fig. 7a-c; Supplementary Fig. 7b). PDPN levels as well as S100A4/PDPN ratio significantly correlated with BRCA1/2 mutational status (Fig. 7b-c). Patients with mutant BRCA1/2 exhibited significantly lower PDPN staining, and higher S100A4/PDPN ratios as compared to BRCA WT patients (Fig. 7b-c). Moreover, multivariate Cox proportional hazards regression analysis of recurrence-free survival, considering S100A4/PDPN ratio and BRCA mutational status showed a strong interaction between the two parameters (Supplementary Table 10). Indeed, when stratified according to BRCA mutational status as well as S100A4/PDPN ratio a striking separation appeared – S100A4/PDPN ratio was a significant classifier of recurrence-free survival in BRCA mutation carriers, but not in BRCA WT patients (Fig. 7d).

**Fig. 7:**
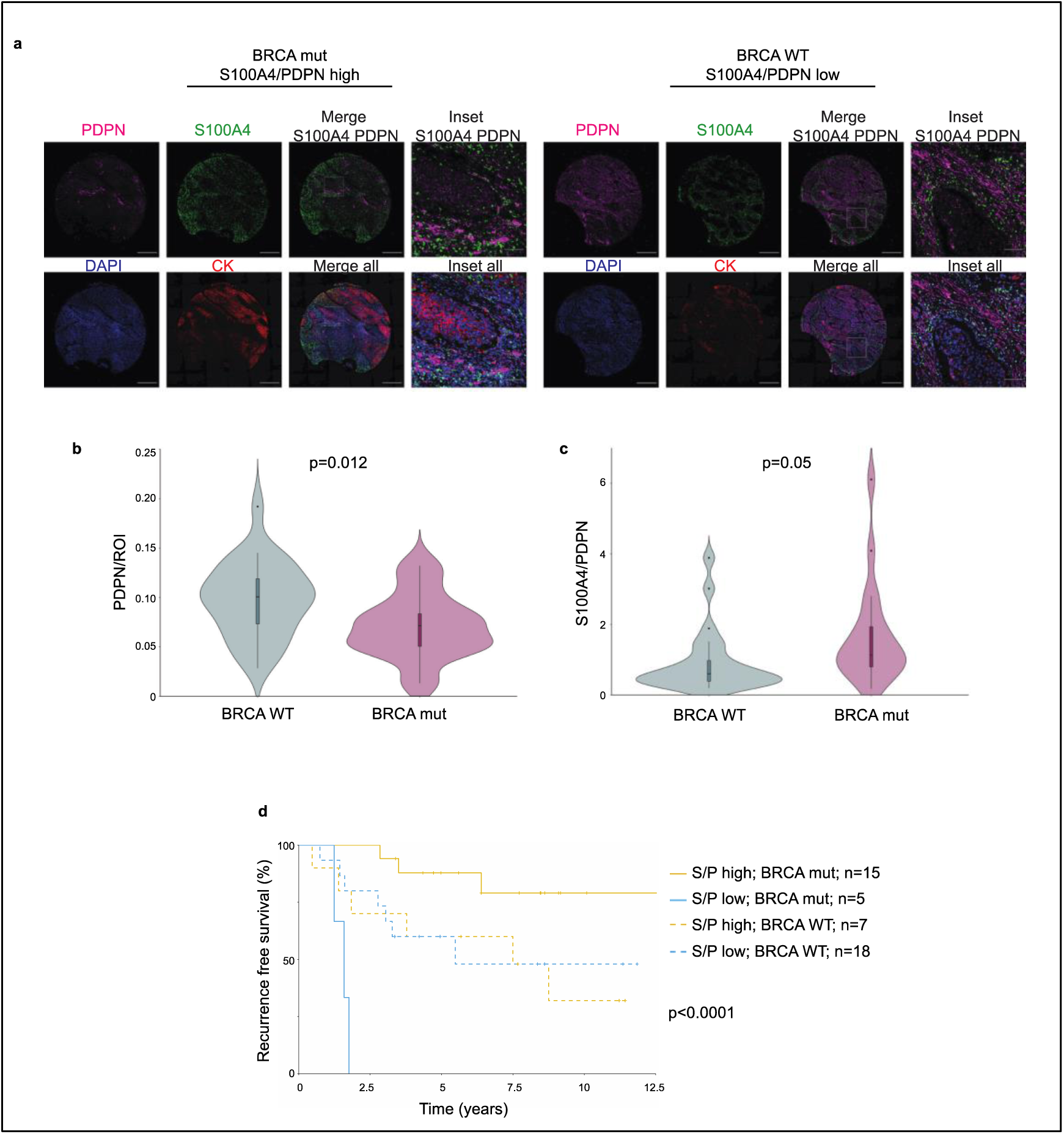
S100A4/PDPN ratio is a classifier of recurrence-free survival in BRCA mutated TNBC. **a,** Representative images of PDPN, S100A4, cytokeratin (CK) and DAPI staining in a BRCA mutated (mut) patient and a BRCA WT patient from our cohort of 72 TNBC patients. Scale bar = 500 μm; inset scale bar = 80 μm **b-c,** Vase-box plots depicting PDPN (**b**) or S100A4/PDPN (**c**) staining scores (see Methods) in BRCA WT (n=25) *vs* BRCA mut (n=20) patients from the TNBC cohort. p Value was calculated using a student’s t-test. **d,** Multivariate analysis through Cox PH model for the TNBC data was performed, then TNBC patients were stratified by BRCA mutational status, and the association of S100A4/PDPN scores (higher vs lower than median) with recurrence free survival was assessed by KM analysis. P-value for the model was calculated using log rank test.

## Discussion

Intratumor heterogeneity is a critical driver of tumor evolution and the main source of therapeutic resistance^18, 27, 28^. Our understanding of how the tumor microenvironment, and in particular CAFs contribute to this heterogeneity is still lacking. Here we find that breast CAFs are comprised of diverse subpopulations that change dynamically over the course of tumor growth and metastasis. These subpopulations cluster into two prototype CAF subtypes, which we term pCAF and sCAF, based on mutually exclusive expression of PDPN in pCAFs and S100A4 in sCAFs. Establishing the relevance of our experimental findings to human disease, pCAFs and sCAFs are major constituents of human breast cancer stroma, and the ratio of S100A4/PDPN expression is a classifier of disease outcome in two independent cohorts of breast cancer.

Recent studies have utilized RNA-sequencing approaches to characterize the tumor microenvironment in different types of cancer^10, 14, 29–32^. In pancreatic cancer, two spatially separated and reversible subtypes of CAFs have been identified by bulk RNA-seq -myofibroblasts (myCAF), located immediately adjacent to the cancer cells, and inflammatory fibroblasts (iCAF), located further away within the dense pancreatic tumor stroma^30^. In breast cancer, a population of matrix remodeling CAFs similar to myCAF was identified and termed mCAFs^14^. Recently, a third population of pancreatic CAFs, antigen-presenting CAFs (apCAFs) was identified by scRNA-seq^31^. Both myCAF and iCAF share similarities with subpopulations of the pCAFs that we have identified, albeit not with sCAFs. In particular, iCAF share common genes with the inflammatory subpopulations of pCAF (*Cxcl1, Il6*), and myCAF are similar to the wound healing pCAF (*Acta2*). In agreement with our analysis suggesting that pCAF originate from tissue resident fibroblasts, both myCAF and iCAF can be derived from tissue resident pancreatic stellate cells. apCAFs, on the other hand, share common genes with sCAFs, in particular with the antigen-presenting sCAFs (*H2-Ab1, CD74, Slpi*), suggesting that these CAFs may serve similar roles in the different tumor types^31^.

In breast cancer, CAFs were recently classified into 4 subclasses with different spatial localization based on a predefined set of cell-surface markers^9^. CAF-S3 in that report were S100A4^High^αSMA^low^, and localized away from cancer cells, as opposed to CAF-S4 which were S100A4^Low^αSMA^High^ and localized closer to cancer cells^9^. The S100A4^High^αSMA^low^ CAFs in that study were not RNA-sequenced or molecularly analyzed, and therefore it is difficult to assess similarities or differences between those CAFs and the sCAF that we have found. However, the localization further away from cancer cells (compared to S100A4^Low^αSMA^High^ CAF) may suggest that these are similar subclasses of CAFs. Another report identified S100A4^High^αSMA^low^ CAFs originating from tissue resident adipocytes^12^. While those CAFs do not share a common morphology or common molecular characteristics with the sCAFs we have identified, both reports highlight the possibility that CAFs originate from cells other than tissue resident fibroblasts.

Indeed, CAFs have heterogeneous origins^10–14^. Three distinct computational approaches (metacell, slingshot and ParTI) point to NMF as the most probable origin of pCAFs. The origin of sCAFs is less clear. While we cannot rule out the possibility that sCAFs originate from NMFs, their transcriptional makeup is disconnected, suggesting that this is not the likely origin of sCAFs. It is also unlikely that sCAFs originated from cancer cells that have undergone EMT, though a minority of cancer cells may have escaped through the negative selection sequencing approach. Rather, we postulate that sCAFs originate from a different mesenchymal source, perhaps from the bone marrow (BM). BM-derived mesenchymal stromal cells (MSC) can be recruited to the tumor and differentiate into CAFs^10, 33, 34^. sCAFs are enriched for several classic MSC markers. Moreover, the molecular chaperone clusterin *(Clu),* recently reported to play a tumor-promoting role in BM-MSC derived CAFs^10^ is among the most differentially upregulated genes in sCAFs (compared to pCAFs). These findings support the hypothesis that sCAFs are derived from MSCs that are recruited to the tumor and differentiate into CAFs. In the tumor sCAFs dynamically shift between several transcriptional programs: cell division, protein translation, and adhesion are the main programs expressed in early tumors. As tumors progress, sCAF are dynamically rewired, and at 4W a subpopulation expressing MHC class II antigen presentation genes takes dominance. MHC class II molecules are constitutively expressed on professional antigen presenting cells (APC). In other cell types, including fibroblasts, the expression of MHC class II can be induced by stimuli such as IFN-γ^35, 36^. Such activation has been shown, for example, in synovial fibroblasts in inflamed joints of rheumatoid arthritis^37^. Antigen presenting CAFs recently described in pancreatic cancer^31^ did not express co-stimulatory molecules. Similarly, we could not detect expression of co-stimulatory molecules in sCAFs. If and how activation of MHC class II in non-professional APC such as the sCAF affects immune responses will be the subject of future investigation.

Metastatic CAFs are poorly defined. In our mouse model, primary tumor CAFs and metastatic CAFs clustered together into the main CAF subtypes pCAFs and sCAFs, suggesting that both subtypes are present in the primary site and in the metastatic site. Nevertheless, primary and metastatic CAFs exhibited distinct subpopulations within each subtype, in particular within pCAFs. These changes are probably driven, to some extent, by the different environment in the lung compared to the mammary tissue. Given the observed shift between 2W and 4W primary tumor CAFs, however, our results suggest that these changes reflect the dynamic rewiring of CAFs along tumor progression, beginning at the primary site and continuing as tumors evolve and metastasize.

The co-existence of different CAF populations, and their dynamic rewiring has both prognostic and potentially therapeutic implications. In two independent cohorts of patients, encompassing together all subtypes of breast cancer, those who had higher sCAF to pCAF ratios had markedly improved survival. In one of these cohorts, comprised only of TNBC patients, high ratios of sCAF/pCAF correlated not only with survival but also with BRCA mutations. BRCA mutations frequently lead to TNBC, and the DNA damage associated with these mutations leads to increased somatic mutational load, and higher T-cell infiltration^26, 38^. It is plausible that the immune regulatory activity of pCAFs inhibits T-cell activation whereas the antigen-presenting sCAFs activate components of the immune system, leading to improved clinical outcome. Our findings highlight the need to define and target deleterious CAF subpopulations, while enriching for potentially beneficial populations, within patient cohorts with defined genetic and transcriptional landscapes.

## Methods

### Ethics statement

All clinical data were collected following approval by the Sheba Medical Center Institutional Review Board (IRB; protocol # 8736-11-SMC) or Ministry of Health (MOH) IRB approval for the Israel National Biobank for Research (MIDGAM; protocol # 130-2013) or as detailed previously^39^. All animal studies were conducted in accordance with the regulations formulated by the Institutional Animal Care and Use Committee (IACUC; protocol # 40471217-2; 09720119-1).

### Human patient samples

Whole tumor sections from 5 ER^+^ and 6 TN breast cancer patients were retrieved and obtained from the Israel National Biobank for Research (MIDGAM; https://www.midgam.org.il/) under MOH IRB# 130-2013 and IRB # 8736-11-SMC. These samples were collected from patients who provided informed consent for collection, storage, and distribution of samples and data for use in future research studies. A tissue microarray (TMA) containing cores from 72 TNBC patients (3 cores per patient), with matching H&Es, was retrieved from the archives of Sheba Medical Center under IRB # 8736-11-SMC. For the METABRIC TMA, appropriate ethical approval from the institutional review board was obtained for the use of biospecimens with linked pseudo-anonymized clinical data as detailed previously^39^.

### Mice

Wild-type BALB/c mice were purchased from Harlan Laboratories and maintained under specific-pathogen-free conditions at the Weizmann Institute’s animal facility.

### Cancer Cell line

4T1 mouse mammary carcinoma cells stably expressing firefly luciferase (pLVX-Luc) were kindly provided by Dr. Zvi Granot, Hebrew University of Jerusalem. For generation of GFP expressing 4T1 cells the FUW-GFP vector was used. GFP expressing cells were selected by two consecutive cycles of FACS sorting. 4T1 cells were cultured in Dulbecco’s modified Eagle’s medium (DMEM) (Biological industries, 01-052-1A) supplemented with 10% fetal bovine serum (FBS) (Invitrogen) in 10-cm tissue culture dishes. For orthotropic injection into the mammary fat pad, the cells were suspended at 80-90% confluence by treatment with 0.25% trypsin, 0.02% EDTA and washed with 1x phosphate buffered saline (PBS).

### Orthotopic injection to the mammary fat pad

BALB/c females were injected at the age of eight weeks, under anaesthesia, with 100,000 4T1-luc cells suspended in 50 μl ice cold PBS, directly into the lower left mammary fat pad. Palpable breast tumors were evident after ten days.

### Normal mammary fat pad isolation and dissociation

Immediately after sacrificing healthy, non tumor-bearing BALB/c females (8W old for single cell analysis, 12W old for bulk sequencing), midline cut was performed and six mammary fat pads were harvested from each animal. For single cell dissociation, tissue was minced using scissors and placed in gentleMACS C tube, where it was disrupted for 25 min at 37°C using gentleMACS dissociator, in the presence of enzymatic digestion solution containing 1 mg ml^-1^ collagenase II (Merck Millipore, 234155), 1 mg ml^-1^collagenase IV (Merck Millipore, C4-22) and 70 unit ml^-1^ DNase (Invitrogen, 18047019) in DMEM (Biological industries, 01-052-1A). The samples were then filtered through a 70 μm cell strainer into ice cold MACS buffer (PBS [-]Ca [-]Mg supplemented with 0.2mM EDTA pH8 and 0.5% BSA) and cells were pelleted by centrifugation at 350g, 5min, 4°C.

### Primary tumor isolation and dissociation

Fourteen or twenty-eight days after 4T1-luc injection, animals were sacrificed and tumors were immediately excised post-mortem. For dissociation to single cells the tissue was minced using scissors and incubated with enzymatic digestion solution containing 3 mg ml^-1^ collagenase A (Sigma Aldrich, 11088793001) and 70 unit ml^-1^ DNase in RPMI 1640 (Biological industries, 01-100-1A) for 20 min at 37°C, with pipetting every 3 min. The samples were then filtered through a 70 μm cell strainer into ice cold MACS buffer and cells were pelleted by centrifugation at 350g, 5min, 4°C. For the enrichment of stromal cells in the samples, single cell suspensions were incubated with 1:1 mixture of anti-EpCAM (Miltenyi, 130-105-958) and anti-CD45 (Miltenyi, 130-052-301) magnetic beads, 20 μl beads per 10^7^ cells 80 μl^-1^, for 15 min at 4°C. The cells were then washed, re-suspended in cold MACS buffer and transferred to LS columns (Miltenyi, 130-042-401) placed on The QuadroMACS separator (Miltenyi, 130-090-976). The column was washed with ice cold MACS buffer and the stromal enriched (CD45, EpCAM depleted) flow-through was collected and pelleted by centrifugation at 350g, 5min, 4°C.

### Lung metastases isolation and dissociation

To allow the growth of >1mm lung metastases, primary tumors were surgically removed under anaesthesia two weeks after injection of 4T1-luc cells to the mammary fat pad.

Following the removal of primary tumors mice were imaged every 4-6 days by in vivo imaging system (IVIS) for detection of luciferase-positive lung metastases, 10 min after intra-peritoneal injection of 225 mg/kg D-luciferin. 2-3 weeks post primary tumor removal the animals were sacrificed and metastases bearing lungs were immediately excised. Metastases were isolated from the lungs and placed in gentleMACS C tubes with an enzymatic digestion solution containing collagenase A 1.5 mg ml^-1^, dispase II 2.5 unit ml^-1^ (Sigma Aldrich, D4693) and DNase I 70 unit ml^-1^ in RPMI 1640. The tissue was disrupted for 30 min at 37°C using gentleMACS dissociator and then the cells were washed with ice cold MACS buffer, filtered through a 70 μm cell strainer and pelleted by centrifugation at 350g, 5min, 4°C.

### Flow cytometry and sorting

Staining was performed on single cells suspended in ice-cold MACS buffer. The antibodies detailed in Supplementary Table 11 were added and the cells were incubated for 30 min on ice. Following staining the cells were washed and resuspended in ice-cold MACS buffer. Single-stain controls were used for compensation of spectral overlap between fluorescent dyes. Propidium idodie (PI) was added shortly before samples were sorted. Cells were sorted with a BD FACSAria Fusion machine and data was analysed using FlowJo software (Tree Star Inc.)

### Single-cell index sorting

Stained cells were single-cell-sorted as previously described^15^. Briefly, cells were sorted into 384-well barcoded capture plates containing 2 µl of lysis solution and barcoded poly(T) reverse-transcription (RT) primers for scRNA-seq^15^. The FACS Diva v8 ‘index sorting’ function was activated to record marker levels of each single cell, and the intensities of all FACS markers were recorded and linked to each cell’s position within the 384-well plate^40^. Four empty wells per 384-well plate were kept as a no-cell control for data analysis. Plates were spun down immediately after sorting to ensure cell immersion into the lysis solution. The plates were then snap frozen on dry ice and stored at −80°C until processing.

### Library preparation for single-cell RNA sequencing

Single-cell MARS-seq libraries were prepared as previously described^15^. In brief, mRNA from cells sorted into MARS-seq capture plates was barcoded, converted into cDNA, and pooled using an automated pipeline. Pooled samples were linearly amplified by T7 in vitro transcription and the resulting aRNA fragmented and converted into a sequencing-ready library by tagging with pool barcodes and Illumina adapter sequences during ligation, reverse transcription and PCR. Library quality and concentration were assessed as described^15^.

### Low-level processing and filtering

All RNA-Seq libraries were sequenced using Illumina NextSeq 500 at median sequencing depth of 28114 reads per single cell. Sequences were mapped to the mouse genome (mm10), demultiplexed, and filtered as previously described^15^, extracting a set of unique molecular identifiers (UMI) that define distinct transcripts in single cells for further processing. We estimated the level of spurious UMIs in the data using statistics on empty MARS-seq wells median noise (2.6%). Mapping of reads was done using HISAT (version 0.1.6;^41^); reads with multiple mapping positions were excluded. Reads were associated with genes if they were mapped to an exon, using the UCSC genome browser as reference. Exons of different genes that shared genomic position on the same strand were considered a single gene with a concatenated gene symbol. Cells with less than 1000 UMIs were discarded from the analysis. After filtering, cells contained a median of 2733 unique molecules per cell. All downstream analysis was performed in R (version 3.6.0).

### Data processing and clustering

The Meta-cell pipeline^42^ was used to derive informative genes and compute cell-to-cell similarity, to compute K-nn graph covers and derive distribution of RNA in cohesive groups of cells (or meta-cells), and to derive strongly separated clusters using bootstrap analysis and computation of graph covers on resampled data. Default parameters were used unless otherwise stated.

Clustering was performed on the CD45^-^ EpCAM^-^ (Supplementary Fig. 1a) compartment of fifteen samples. Cells with high expression of *Hbb-b1* or *Ptprc* were regarded as contaminants of red blood or immune cells respectively, and were discarded from subsequent analysis. Following clustering of the remaining cells (Supplementary Fig. 1d), cells with high expression of *Pecam1* and *Rgs5* were identified as endothelial cells and pericytes respectively, and discarded from further analysis. In addition, a group of 33 cells with markedly high expression of *Mki67* and *Myc* was assumed a contamination of cancer cells and removed from further analysis.

Meta-cell clustering was performed over the top 10% most variable genes (high var/mean), with total expression over 50 UMI and more than 2 UMI in at least three cells, resulting in 1017 feature genes. After clustering, resulting clusters were filtered for outliers, and cells with more than 4 fold deviation in expression of at least one gene were marked as outliers and discarded from further analysis. This resulted in 43 outlier cells and retained 8033 cells for further analysis.

In order to annotate the resulting meta-cells into cell types, we used the metric FPgene,mc, which signifies for each gene and meta-cell the fold change between the geometric mean of this gene within the meta-cell and the median geometric mean across all meta-cells. We used this metric to “color” meta-cells for the expression of subset specific genes such as *Gsn* and *S100a4*. Each gene was given a FP threshold and a priority index. The selected genes, priority, and fold change threshold parameters are as detailed in Supplementary Table 12.

### GO enrichment analysis

Gene set enrichment analysis was done using Metascape software (http://metascape.org).

### Trajectory finding

To infer trajectories and align cells along developmental pseudotime, we used the published package Slingshot^19^. In short, Slingshot is a tool that uses pre-existing clusters to infer lineage hierarchies (based on minimal spanning tree, MST) and align cells in each cluster on a pseudotime trajectory. We applied Slingshot on the pCAF population of the primary tumor (2W, 4W). We chose a set of differential genes between the clusters (FDR corrected chi-square test, q < 10-3, fold change > 2). We performed PCA on the log transformed UMI, normalized to cell size. We ran Slingshot on the top five principal components, with *Pdpn^+^* cells from the normal mammary fat pads as the starting cluster.

### Pareto analysis

#### sCAF Single-Cell Data

The gene expression dataset of NMF, 2W and 4W CAFs included 21948 genes and 6587 cells (3067 sCAFs and 3521 NMF & pCAFs). In PCA analysis of the sCAFs, cells from 2W and 4W timepoints formed a continuum whereas the Mets formed a separate cluster. We therefore excluded the Mets from ParTI analysis, which focuses on continuous expression patterns. We considered cells with a total of at least 3×10^3^ UMIs and genes with at least 10^3^ UMIs, totaling 2292 cells and 790 genes. Each cell was down-sampled to 10^3^ UMIs, and each gene was log transformed and centered by subtracting its mean.

### Data Dimensionality

To determine the dimensionality of the data for ParTI, we used PCHA to find the best-fit polytopes with k=3-7 vertices (k=3 is a 2D triangle, k=4 is a 3D tetrahedron, and so on). We calculated the variance of the vertex positions by PCHA on bootstrapped data (resampling the cells with returns). We found that the variance for k=3 and 4 is low, and rises sharply for k>4 (Supplementary Fig. 3f), indicating that it is not possible to determine the positions of more than 4 vertices with high reliability. In agreement with the three-dimensionality of the tetrahedron, PCA analysis indicated that the first 3 PCs explain much more variance than higher order PCs (Supplementary Fig. 3e). We concluded that 4 vertices and the first three principal components are the appropriate choice for this analysis.

### Tetrahedron Significance

The variation in the vertex positions of the real data (bootstrapping) was much smaller than the variation of the vertex positions in the best-fit tetrahedron (PCHA) for 1000 shuffled datasets (p<0.001; Supplementary Fig. 3g-h). We further tested the statistical significance of the tetrahedron by the t-ratio test as described in^21, 43^. The t-ratio is the ratio between the convex hull of the data and the minimal enclosing tetrahedron. A large t-ratio means that the tetrahedron hugs the data tightly. The observed t-ratio was significantly larger than the t-ratios of shuffled datasets (p=6×10^−3^, Supplementary Fig. 3i).

### Enrichment Calculation and GO analysis

We defined enriched genes for each vertex by calculating the Spearman rank correlation between the gene’s expression and the Euclidean distance of cells from the vertex in gene expression space. We call a gene enriched if its expression shows a correlation coefficient below −0.2 with a statistically-significant p-value controlled for multi-hypothesis testing by a false discovery rate (FDR) correction using the Bonferroni procedure with a threshold of 10^-5^. GO analysis was performed using MathIOmica^44^ with a cutoff of at least 3 genes for each GO term. To address circularity concerns stemming from using gene expression both to infer the position of the vertex and their functions, we use a leave-one-out procedure: for each enriched gene, we recompute the position of the vertices after removing the gene. We then determine which samples are closest to the new vertices, and test whether the gene is still significantly enriched close to the vertex by the same method as above.

### Vertex temporal ordering

We computed the relative representation of cells from each time point (2W, 4W) between the 4 vertices (Fig. 3d). Cells from each time point were down sampled to reach the same number (300), and the fraction of each time point in the 100 cells closest to each archetype was calculated. Error bars were calculated by bootstrapping (10^3^).

### Bulk RNA sequencing

10^4^ cells were sorted from each population using the gating strategy described for single cell RNA-seq, with the addition of PDPN as a positive selection marker for pCAFs (Supplementary Fig. 1a). NMF were taken from the CD45-/EpCAM-population without further selection. sCAFs were collected based on negative selection for all markers (CD45^-^EpCAM^-^PDPN^-^). The cells were collected directly into 40 µl of lysis/binding buffer (Life Technologies) and mRNA was isolated using 15 μl dynabeads oligo (dT) (Life Technologies). mRNA was washed and eluted at 70°C with 6.5 μl of 10 mM Tris-Cl pH 7.5. RNA-seq was performed as previously described^15^. Libraries were sequenced on an Illumina NextSeq 500 machine and reads were aligned to the mouse reference genome (mm10) using STAR v2.4.2a^45^. Duplicate reads were filtered if they aligned to the same base and had identical UMIs. Read count was performed with HTSeq-count^46^ in union mode and counts were normalized using DEseq2^47^.

### Tracing of host *vs* cancer markers in sCAFs

To test for presence of 4T1 cells in the negatively selected sCAF population we traced the LTR of a luciferase plasmid expressed in these cells. While this sequence could not be detected by scRNA-seq (due to polyA selection) we could detect it by bulk RNA-seq. We therefore counted the number of reads mapped to the LTR in different populations from bulk FACS sort and normalized these to the number of reads mapped to the house keeping gene Actin. The normalized LTR reads were 44-fold more abundant in bulk EPCAM^+^ cells from the tumor (that may also contain normal epithelial mammary cells) than in sCAFs (0.011 in EPCAM^+^ *vs* 0.00027 in sCAFs). We could not detect LTR reads in pCAFs and NMFs. These results suggest that the majority of sCAF do not originate from 4T1 cancer cells, however there is a low level of contamination by 4T1 cells, at least in the bulk population.

To validate these results we expressed GFP in 4T1 cells, injected these into the mammary fat pad of mice, dissociated the resulting tumors, FACS sorted to remove PDPN^+^ and CD45^+^ cells, and then further sorted for bulk RNA-seq of the following populations: (1) GFP^+^EpCAM^+^ (expected to include 4T1 cells); (2) GFP^+^EpCAM^-^ (expected to include 4T1 cells that may have undergone EMT); (3) GFP^-^EpCAM^+^ (expected to include host epithelial cells); and (4) GFP^-^EpCAM^-^ (expected to include sCAF, as well as a minor population of endothelial cells and pericytes). Bulk sequencing followed by differential gene expression analysis confirmed that GFP^+^EpCAM^+^, GFP^+^EpCAM^-^, and GFP^-^EpCAM^+^ populations exhibited similar patterns of gene expression, whereas GFP^-^EpCAM^-^ cells were distinct, suggesting that GFP^-^EpCAM^-^PDPN^-^CD45^-^ cells do not originate from cancer cells, nor do they resemble normal epithelial cells (Supplementary Table 5). The top 20 differentially upregulated genes in GFP^-^EpCAM^-^ cells (i.e. sCAFs) compared to GFP^+^EpCAM^+^ cells (i.e. 4T1 cells) contain classic stromal genes such as *Cxcl12*, *Col3a1* and *Ccl4* (Supplementary Table 5). The classic epithelial marker *Krt14* is among the most differentially downregulated genes in GFP^-^EpCAM^-^ cells compared to GFP^+^EpCAM^+^ cells (Supplementary Table 5).

### FACS sorting for functional assays with pCAFs

Primary tumors were harvested and dissociated into single cell suspension four weeks after injection of 4T1-luc cells as described above. Pelleted cells were re-suspend in 5 ml RBC lysis buffer (BioLegend 420301) for 10 min in RT and washed with 20 ml MACS buffer, afterwhich depletion of CD45 and EpCAM was performed as described above. For pCAF enrichment, the CD45, EpCAM depleted fraction was incubated with PDPN-biotin antibody for 30 min on ice, the cells were washed and incubated with anti-biotin magnetic beads (Miltenyi, 130-090-485), 20 μl beads per 10^7^ cells 80 μl^-1^, for 15 min at 4°C. The PDPN enriched cell suspension was isolated using LS columns as described above, and the cells were stained for Ter119-PB, CD45-BV711, EpCAM-AF488, PDPN-APC, and Ly6C-PerCP/Cy5.5. PDPN^+^ cells were gated as described in Supplementary Fig. 1a. The PDPN^+^ cells were then sorted into Ly6C^+^ and Ly6C^-^ populations.

### CD8 T cell isolation and CFSE labeling

5*10^4^ Ly6C^+^ or Ly6C^-^ pCAFs were plated in 96-wells in presence of RPMI 1640 supplemented with 10% FBS. Three days later CD8^+^ cells were isolated and added to the Ly6C^+^ or Ly6C^-^ pCAFs. For CD8^+^ T-cells isolation, spleens were harvested post mortem and dissociated into single cell suspensions. Red blood cells were depleted by red-blood lysis buffer. CD8 positive cells were isolated by a positive selection kit (CD8a (Ly-2) Microbeads, mouse, Mitenyi 130-117-044). Cells were then stained with 2μM CFSE and incubated with CD3C/D28 Dynabeads^TM^ for 48-72h either with or without CAFs. For FACS analysis magnetic beads were removed and DR^TM^ (BioLegend) was used for exclusion of dead cells from the analysis. FACS analysis of the CFSE stained CD8^+^ cells was performed using Kaluza software version 2.1 (Beckman Coulter).

### Flow cytometry of pCAFs, sCAF and NMF markers

Tissues were harvested and dissociated into single cell suspensions as described above. Prior to intra cellular staining of pCAF markers, cells were fixed with 4% PFA in PBS for 10 min, washed and resuspended in 200μl permabilization/washing buffer (PBS [-]Ca [-]Mg, 0.1% TWEEN 20 (BIO BASIC), 1% BSA) and incubated for 20 min RT. The permeabilized cells were stained with CD45-BV711, EpCAM-AF488, PDPN-APC, Ly6C-PerCP/cy5.5 and αSMA-FITC. For sCAF marker staining, live cells were stained with CD45-BV711, EpCAM-AF488, PDPN-APC and I-A/I-E-APC/Cy7.

### Immunohistochemistry of mouse tissues

Normal mammary fat pads, tumors or metastases dissected as described above were fixed in 4% Paraformaldehyde (PFA), processed and embedded in paraffin blocks, cut into 4μm sections and immunostained as follows: Formalin-fixed, paraffin-embedded (FFPE) sections were deparaffinized, treated with 1% H2O2 and antigen retrieval was performed by microwave (2-3 min until boiling followed by low power heating for 10 min and then cooling at RT for 10 min) with Tris-EDTA buffer (pH 9.0). Slides were blocked with 10% normal horse serum (Vector Labs, S-2000), and the antibodies listed in Supplementary Table 11 were used. Visualization was achieved with 3,30-diaminobenzidine (DAB) as a chromogen (Vector Labs kit #SK4100). Counterstaining was performed with Mayer-hematoxylin (Sigma-Aldrich MHS-16). Images were taken with a Nikon Eclipse Ci microscope.

### Immunofluorescent staining of mouse and human tissues

Whole FFPE sections from mouse and human tumors, and cores from the human TNBC and METABRIC TMAs, were deparaffinized and then all tissues were incubated in 10% Neutral buffered formalin (NBF prepared by 1:25 dilution of 37% formaldehyde solution in PBS) for 20 min in room temperature, washed (with PBS) and then antigen retrieval was performed by microwave with citrate buffer (pH 6.0; for PDPN) or with Tris-EDTA buffer (pH 9.0; for CK and S100A4). Slides were blocked with 10% BSA + 0.05% Tween20 and the antibodies listed in Supplementary Table 11 were diluted in 2% BSA in 0.05% PBST and used in a multiplexed manner with the OPAL reagents, each one O.N. at 4°C. The OPAL is a stepwise workflow that involves tyramide signal amplification. This enables simultaneous detection of multiple antigens on a single section by producing a fluorescent signal that allows multiplexed immunofluorescence (MxIF), imaging and quantitation (Opal reagent pack and amplification diluent, Perkin Elmer). Briefly, following overnight incubation with primary antibodies, slides were washed with 0.05% PBST, incubated with secondary antibodies conjugated to HRP for 10 min, washed again and incubated with OPAL reagents for 10 min. Slides were then washed and microwaved for 10 min, washed, stained with DAPI and mounted. We used the following staining sequences: CK → S100A4 → PDPN → DAPI (for mouse); S100A4 → CK → PDPN → DAPI (for human); S100A4 → CD45 →DAPI; or CD3 →DAPI.

Whole tumor sections from the human TNBC cohort were stained by either of the following sequences: SMA → CK → PDPN→ DAPI, or S100A4 → NT5E → HLA→ CK → DAPI.

Each antibody was validated and optimized separately, and then multiplexed immunofluorescence (MxIF) was optimized to confirm that signals were not lost or changed due to the multistep protocol. Slides of mouse and whole human tumor sections were imaged with a DMi8 Leica confocal laser-scanning microscope, using either a HC PL APO 20x/0.75 objective, a HC PL APO 40x/1.3 oil-immersion objective or a HC PL APO 60x/1.4 oil-immersion objective and HyD SP GaAsP detectors. TMA slides were imaged with an Eclipse TI-E Nikon inverted microscope, using a CFI supper Plan Fuor 20X/0.45 NA and DAPI/FITC/Cy3 and Cy5 cubes. Images were acquired with cooled electron-multiplying charge-coupled device camera (IXON ULTRA 888; Andor).

### Image analysis

Quantification of TMA staining was done using Fiji image processing platform^48^. Regions of interest (ROIs) were manually depicted to include all intact tissue areas and exclude regions of adipose tissue (due to its tendency to display nonspecific staining). H&Es from the TNBC TMA were used to assist in training and optimizing this step. Following background subtraction using a rolling ball with a radius of 200 pixels, the CK, S100A4, PDPN channels were thresholded using Otsu’s method. The threshold of CD3 (which was stained and analyzed separately) was set to 2500-65535. All pixels above the threshold were counted as 1, and their sum was divided by ROI (Supplementary Fig. 5b). Channel/ROI scores of all replicate cores from the same patient (typically 3) were averaged and the average score was used for statistical analysis as described in the following section. Ratios between different stains were calculated by dividing the channel/ROI scores of each core, and then averaging the scores of each patient. In the TNBC cohort, two patients were excluded from the analysis since their S100A4/PDPN values were 3 standard deviations higher than the average. 4 patients were excluded from CD3 analysis due to unusually high background staining that could not be interpreted. All other scores collected were included in the analyses. In the METABRIC cohort 5 patients were excluded from the analysis since their S100A4/PDPN values were 3 standard deviations higher than the average.

Regional analysis of cancer-adjacent and dense-stromal regions was performed as follows: We applied a threshold to the CK channel using Moments method and expanded the CK^+^ regions using the “Dilate” function six times. A mask generated from this image was used to define “cancer-adjacent” regions, and the inverse mask was used to define “dense-stromal” regions. Ratios between different stains were calculated for each region as described above.

Analysis of overlap between CAF markers in human breast tumors was performed on MxIF staining of whole tissue FFPE sections from the TNBC cohort. Briefly, the sections were scanned by confocal microscope as described above. In cases of staining with more than 4 fluorophores we performed linear spectral unmixing, using a matrix that calculates the overlap between the different fluorophores in cases of overlapping emission wavelengths. We calculated the matrix by imaging each individual fluorophore at the excitation/emission wavelengths of each channel, and subtracted the relative contribution of the overlapping channels from the original image. The images from each channel were then z-stacked and a threshold was applied using Moments method to generate masks. The number of overlapping pixels between channels was quantified using the “AND” function in the image calculator. The number of overlapping pixels was then divided by the total number of pixels of the originating channels.

Analysis of the overlap between CAF markers in murine 4T1-tumors was performed on MxIF staining of whole tissue FFPE sections. Masks for each channel were generated using Moments method. Since CK is located only in the cytoplasm while in the mouse S100A4 is observed, in some cases, also in the nucleus, we removed the nuclear region from each channel prior to the analysis. Briefly, we applied “Fill holes” and “Watershed” on the DAPI mask, removed particles smaller than 8μm^2^ and created a mask from the resulting particles. The “Subtract” command in the image calculator was used to remove the nuclear region from each channel. The number of overlapping pixels between channels was quantified using the “AND” function in the image calculator. The number of overlapping pixels was then divided by the total number of pixels of the originating channels.

### Statistical Analysis

Clinical characteristics were compared by means of the Pearson χ2 test for categorical variables, and a student’s t-test for age (continuous variable). Recurrence free and overall survival rates were obtained based on Kaplan-Meier estimates and a log rank test was performed to study the difference of recurrence free / overall survival rates. Density estimate of the divided values were obtained using integrated vase-box plots, the means of the two genetic groups were compared using a student’s t-test. For visualization purposes, values that were above the two box plot whiskers were omitted from the plot, but included in the statistical analysis (7 values from Supplementary Fig. 6b, 1 value from Supplementary Fig. 6d, and 6 values from Supplementary Fig. 7c). Relative Risk estimates and 95% confidence intervals (CIs) were calculated utilizing Cox proportional hazard regression model for the recurrence free survival data, univariate analysis to study the effects of the variables on recurrence free survival, and multivariate analysis considering first order interaction. Dividing continuous variables: To visualize the results of the Cox proportional hazard regression model, S100A4/PDPN and PDPN/Total ROI were divided into High/Low groups by their median (in the TNBC cohort) each by their median, or by a value of 1 (in the METABRIC cohort).

