## Supplemental Figures and Tables for "Cancer-associated fibroblast compositions change with breast-cancer progression linking S100A4 and PDPN ratios with clinical outcome"

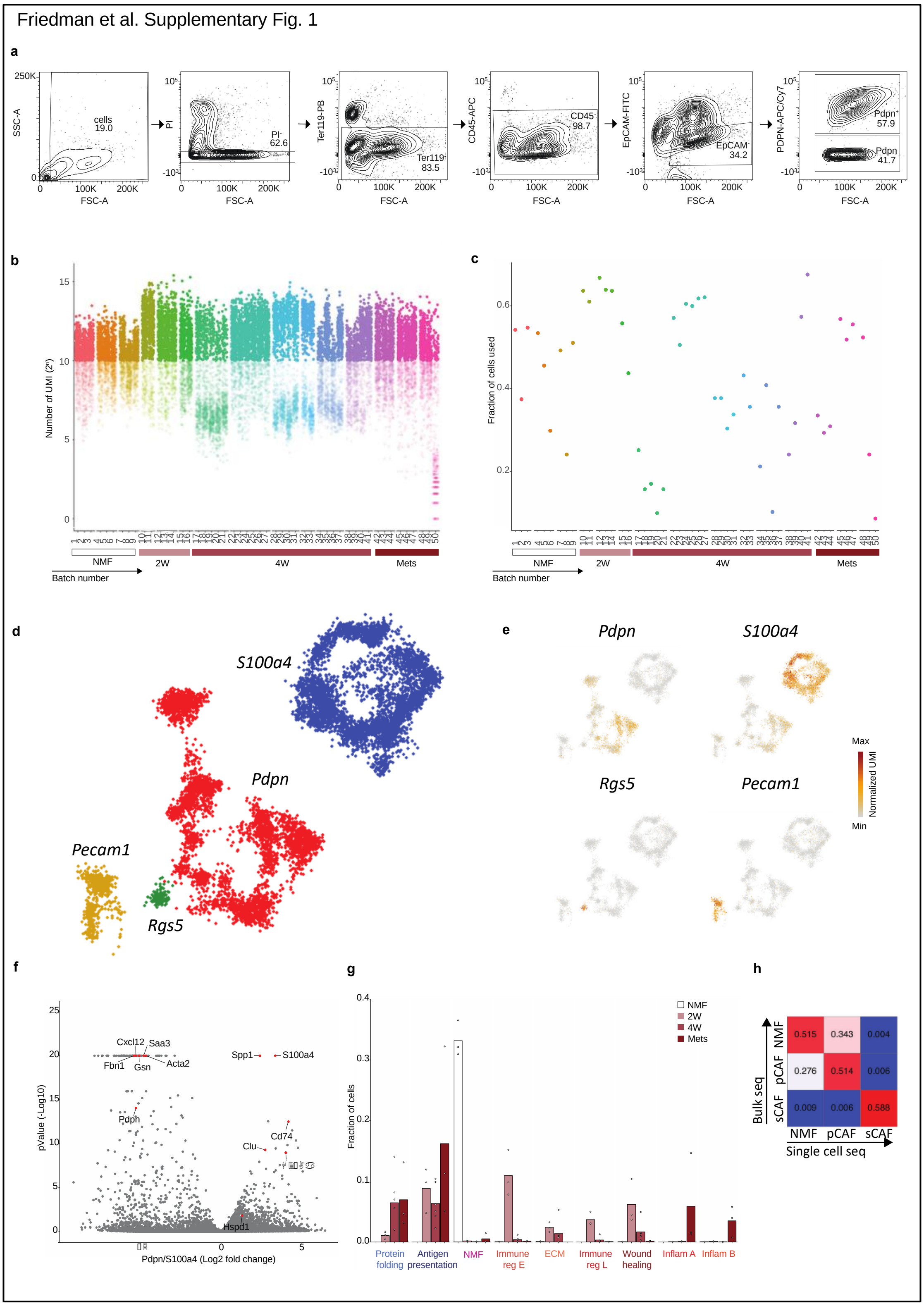


**Supplementary Fig. 1: A single cell map of breast cancer stroma. a,** Sorting strategy: All live single cells (PI negative cells after debris and doublet exclusion) staining negative for Ter119 (Red blood cells); CD45 (immune); and EpCAM (epithelial) were collected and single cell sorted. PDPN was used for index sorting of pCAFs. Representative flow cytometry plots are shown. **b-d,** Quality control metrics of single cells analyzed in this study. **b,** Total unique molecular identifier (UMI) per cell. Cells are grouped by batch (plate) and color-coded by biological replicate (mouse). The time point for each batch is indicated. Cells with less than 1,000 UMI were discarded from the analysis. **c,** Fraction of analyzed cells/batch after filtering. Batches are grouped and color-coded as described in **b**. **d,** Single cell RNA-seq data from 8987 QC positive cells staining negative for Ter119, CD45 and EpCAM was analyzed and clustered using the MetaCell algorithm, resulting in a two-dimensional projection of cells from 15 mice. 88 meta-cells were associated with 4 broad clusters, annotated and marked by color code. **e**, Expression of the hallmark genes for the 4 clusters presented in **d** on top of the two-dimensional projection of breast cancer stroma. Colors indicate log transformed UMI counts normalized to total counts per cell. **f,** Volcano plot displaying differentially expressed genes between *Pdpn*$^{+}$ fibroblasts and *S100a4*$^{+}$ fibroblasts (see also supplementary table 4). Marker genes for NMF, pCAF, and sCAF are highlighted. **g,** Fraction of cells originated from each mouse and subset, from all cells originated in their time point. Bar values represent the mean fraction values. Time points and subclasses are annotated and colored as in Fig. 1d. **h,** Squared Pearson correlation matrix between bulk and single cell sequencing results for NMF, pCAF, and sCAF.

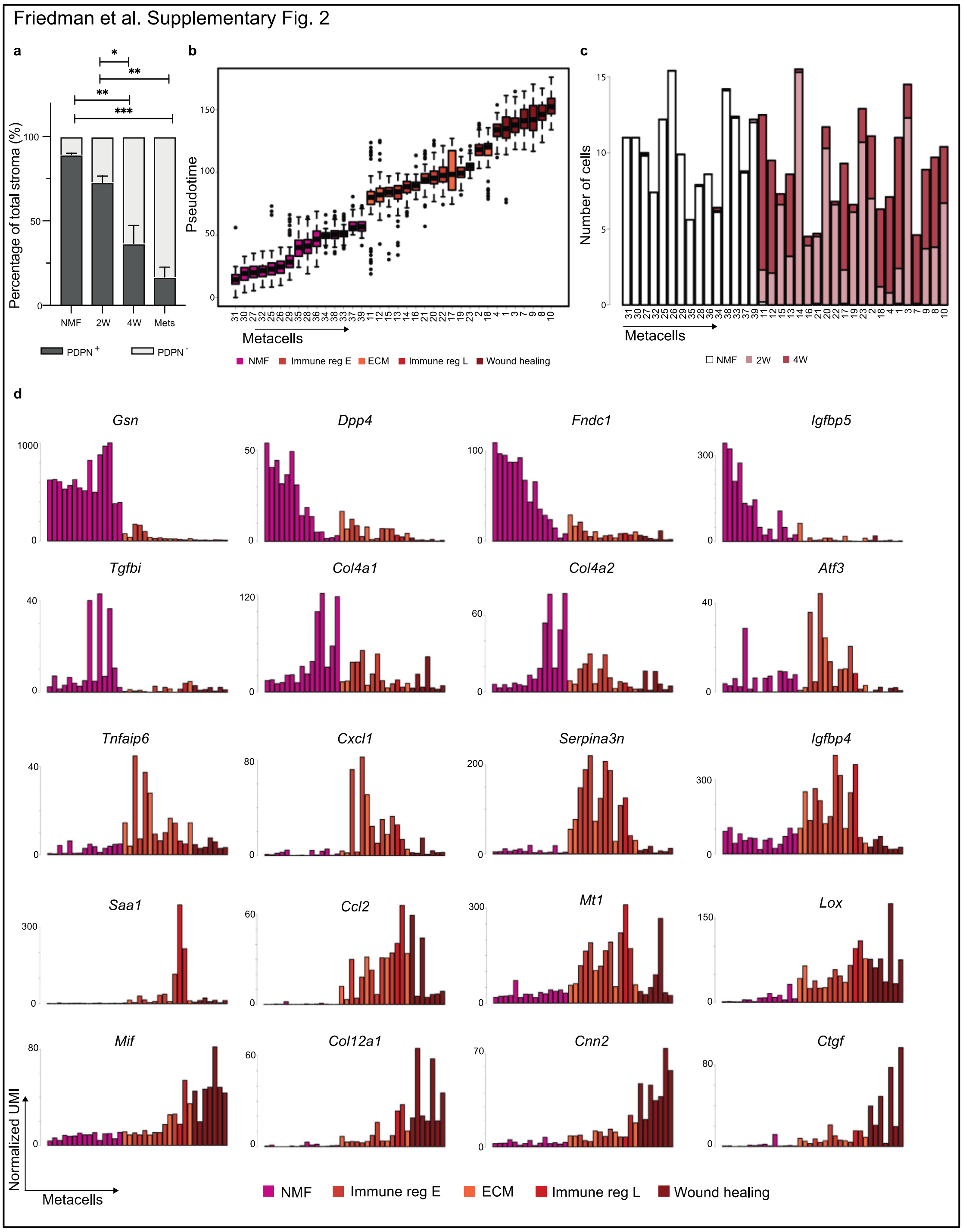


**Supplementary Fig. 2: *Pdpn*+ fibroblasts undergo dynamic changes in gene expression and subset composition during tumor progression. a,** Cell-surface PDPN protein expression levels obtained from index sorting data (presented in Fig. 1d, lower panel), were used to quantify the percent of $PDPN^+$ and $PDPN^-$ cells in the $CD45^-EpCAM^-$ stroma in the different time points. P-value was determined by Anova considering multiple comparisons. **b,** Pseudo-time of expression for individual metacells (color coded by functional subclasses as in Fig. 2) included in the slingshot analysis. Box plots display median bar, first–third quartile box and 5th–95th percentile whiskers. **c,** Distribution of cells across time points (color coded) within metacells included in the slingshot analysis. Metacell numbers and order are consistent across all figure panels and match the order in Fig. 2. **d,** Expression of hallmark

NMF and pCAF genes (additional to those presented in Fig. 2e) across metacells (average UMI/cell), ordered by pseudo-time. * $p<0.05$; ** $p<0.005$; *** $p<0.001$

Friedman et al. Supplementary Fig. 3

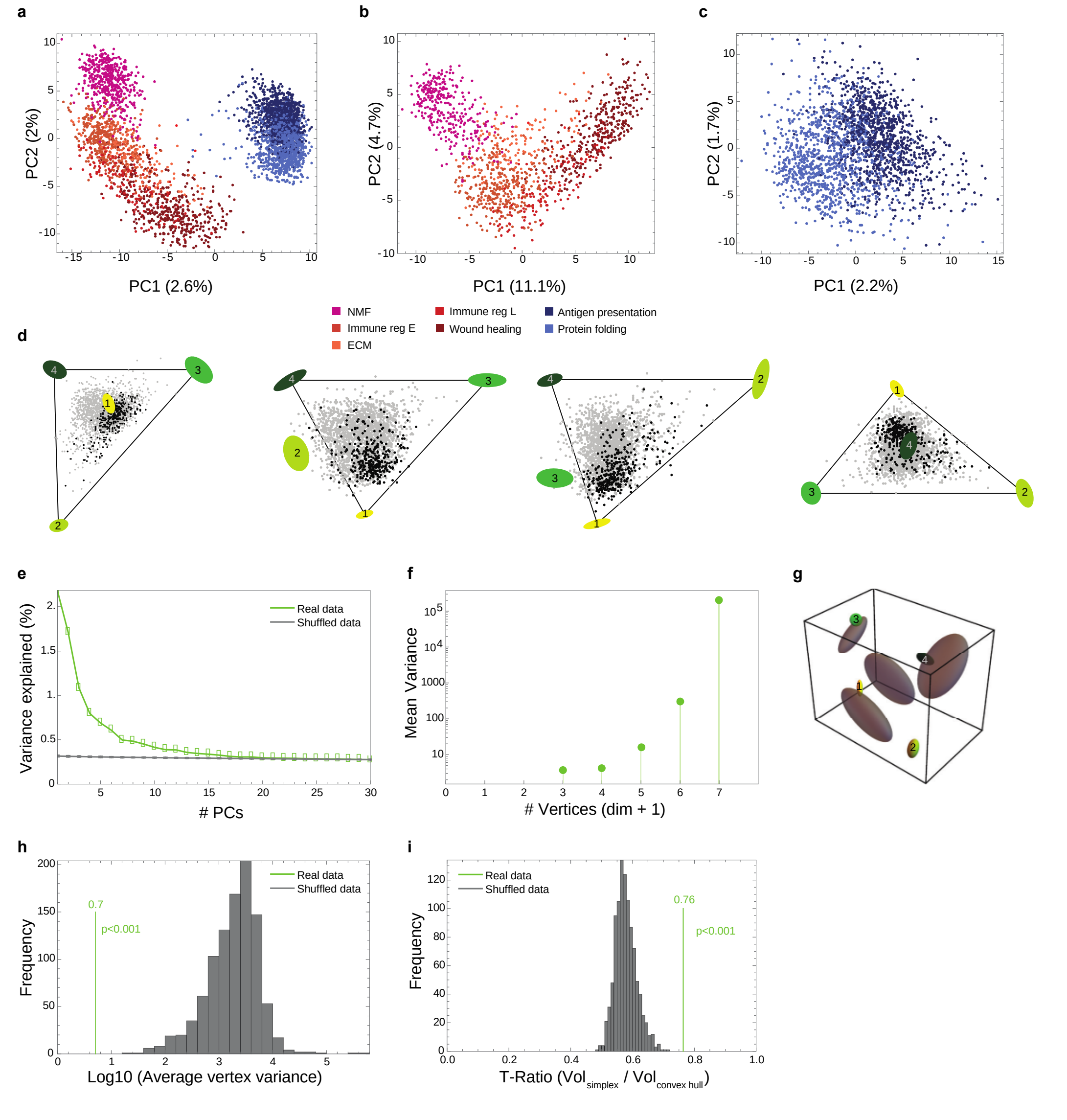


**Supplementary Fig. 3: pCAFs and NMFs form a curve in gene-expression space, whereas a tetrahedron describes sCAF gene expression. a,** PCA analysis of NMF, and pCAF and sCAF from 2W and 4W, color coded according to the subclasses defined in Fig. 1c. **b-c,** PCA analyses for NMF and pCAF (**b**) and for sCAF (**c**) color coded as in **a**. **d,** Data projected on the four faces of the tetrahedron. **e,** Explained variance as a function of the number of PCs (real data) vs. random. Note that the total variance explained by the first 3 PCs, about 5%, is typical of single-cell gene expression data[22]. **f,** Variance of vertex positions as a function of the number of vertices considered, using PCHA with k=3-7 vertices. **g,** Variation of vertex position (bootstrapping) for the real data (ellipses color-coded as in Fig. 3) *vs* shuffled data (grey ellipses). **h,** Histogram depicting the average variation of vertex positions calculated for the real data (green) *vs* multiple runs of shuffled data (grey). **i,** Histogram depicting the ratio between the volumes of the convex hull of the data and the minimal enclosing tetrahedron (t-ratio). The t-ratio of the real data (green) is compared to t-ratios of shuffled data (1000 shuffles; grey).

Friedman et al. Supplementary Fig. 4

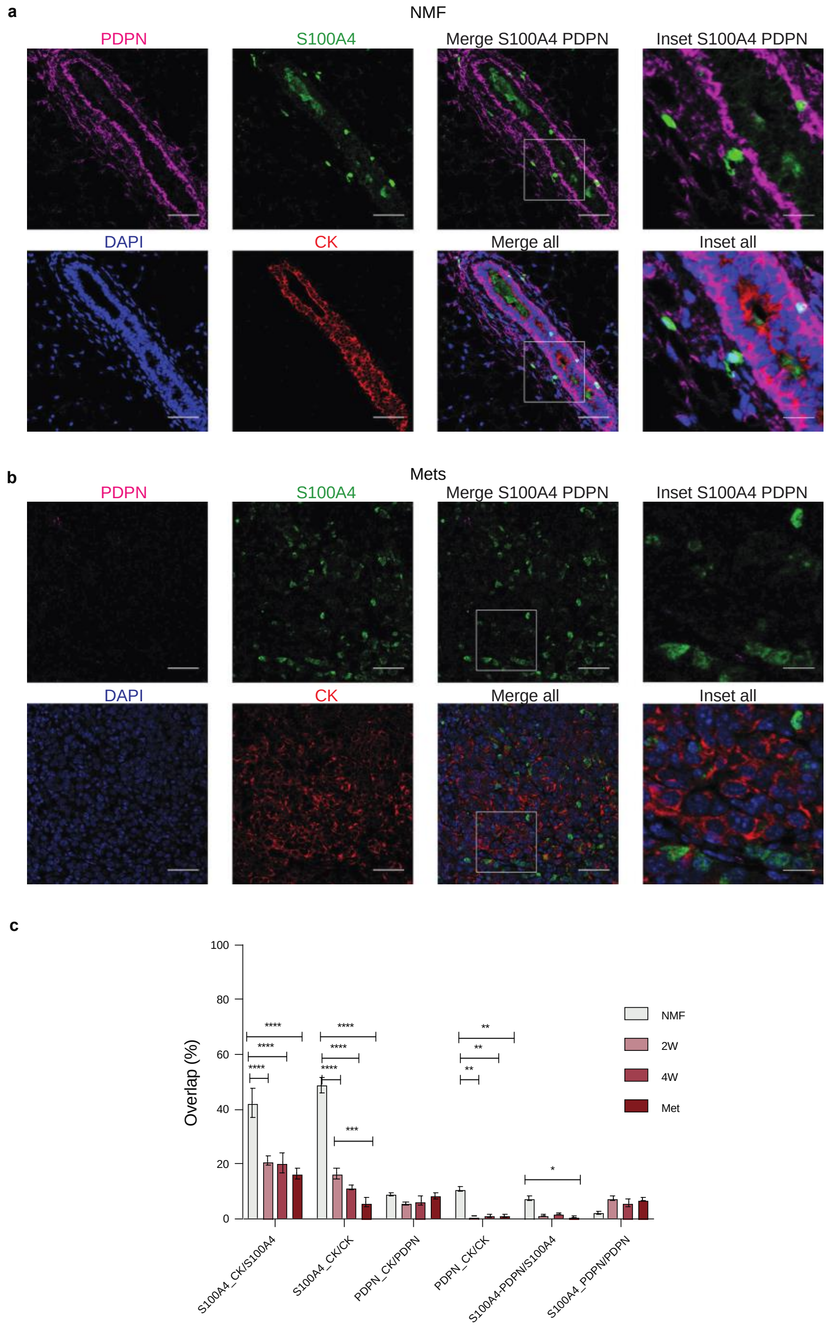


**Supplementary Fig. 4: PDPN and S100A4 proteins mark distinct types of cells in mouse tumors, the majority of which are CK-negative. a-b,** Representative images of normal mammary fat pads (NMF; **a**) and lung metastases (Mets; **b**) (see Fig. 4a) stained with antibodies against the indicated proteins. Scale bar = 50μm, inset scale bar = 17□m. **c,** Quantification of the average overlap between CK, PDPN, and S100A4 staining in NMFs, primary tumors (2W and 4W) and Mets. Bars represent the number of overlapping pixels between two channels, divided by the total number of pixels of the originating channels, averaged for 3 biological replicates (9 images per mouse). P-values were calculated by two-way Anova correcting for multiple comparisons. * p<0.05; ** p<0.005; *** p<0.001; *** p<0.0001

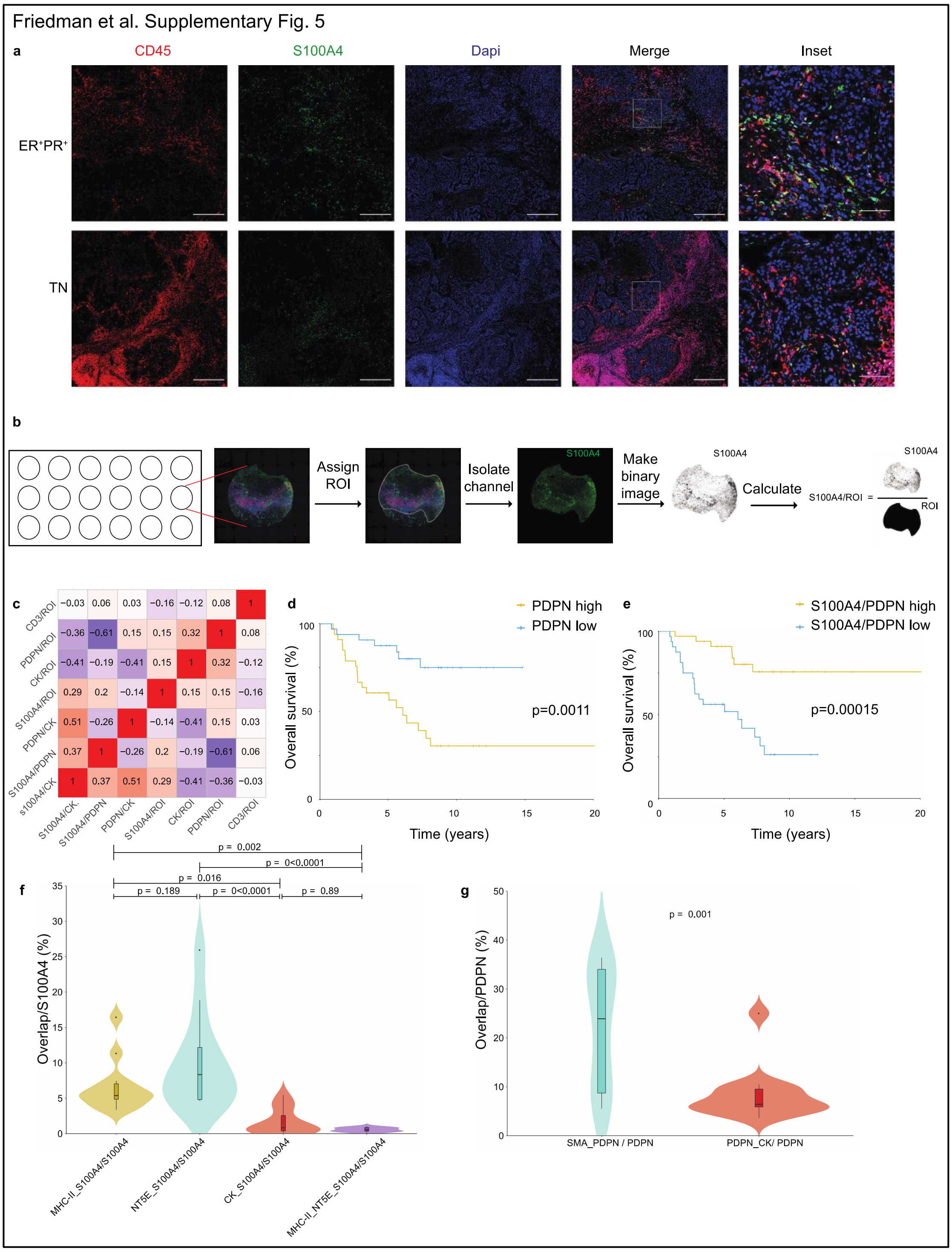


**Supplementary Fig. 5: Low PDPN staining and high S100A4/PDPN staining ratios correlate with better overall survival of TNBC patients. a,** Representative images of MxIF staining of serial sections from the same patients presented in Fig. 6a with antibodies against the indicated proteins. Scale bar = 500 m.; inset scale bar = 90 m **b,** Illustration of the image analysis workflow. **c,** Heat map showing Pearson's correlation coefficients of the staining scores for different cell type markers. **d-e,** The association with overall survival of PDPN (**d**) or S100A4/PDPN (**e**) scored and classified as in Fig. 6b was assessed by KM analysis. p Values were calculated using log rank test. **f-g,** The overlap between S100A4, CK, MHC-II and NT5E stains (**f**) and between PDPN, CK, and SMA stains (**g**) in a subset of

12 patients (average scores of 3 images per patient) of the TNBC is presented. P-values were calculated by two-way Anova (with correction for multiple comparisons in (**f**)).

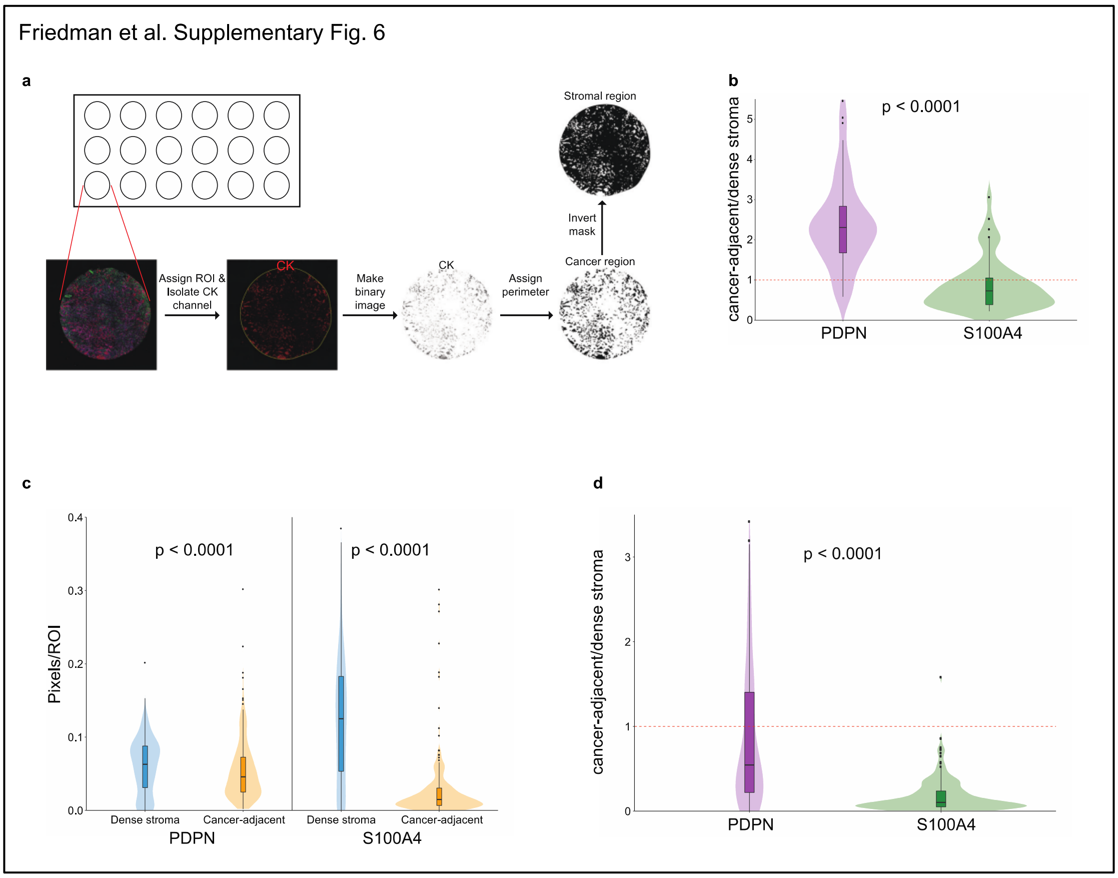


**Supplementary Fig. 6: pCAFs tend to localize to cancer-adjacent regions more often than sCAFs in human breast cancer patients. a,** Illustration of the regional analysis workflow. **b,** The ratio of cancer-adjacent/dense stroma PDPN and S100A4 staining was determined for each core in the TNBC TMA (See also Fig. 6d). P-value was calculated using Wilcoxon matched pairs signed rank test. **c-d,** Cancer-adjacent regions and regions of dense stroma were determined for each core in the METABRIC TMA based on CK staining (see Methods section), PDPN and S100A4 staining in each region was scored (**c**) and the ratio of cancer-adjacent/dense stroma PDPN and S100A4 staining was determined (**d**). P-values were calculated using Wilcoxon matched pairs signed rank test.

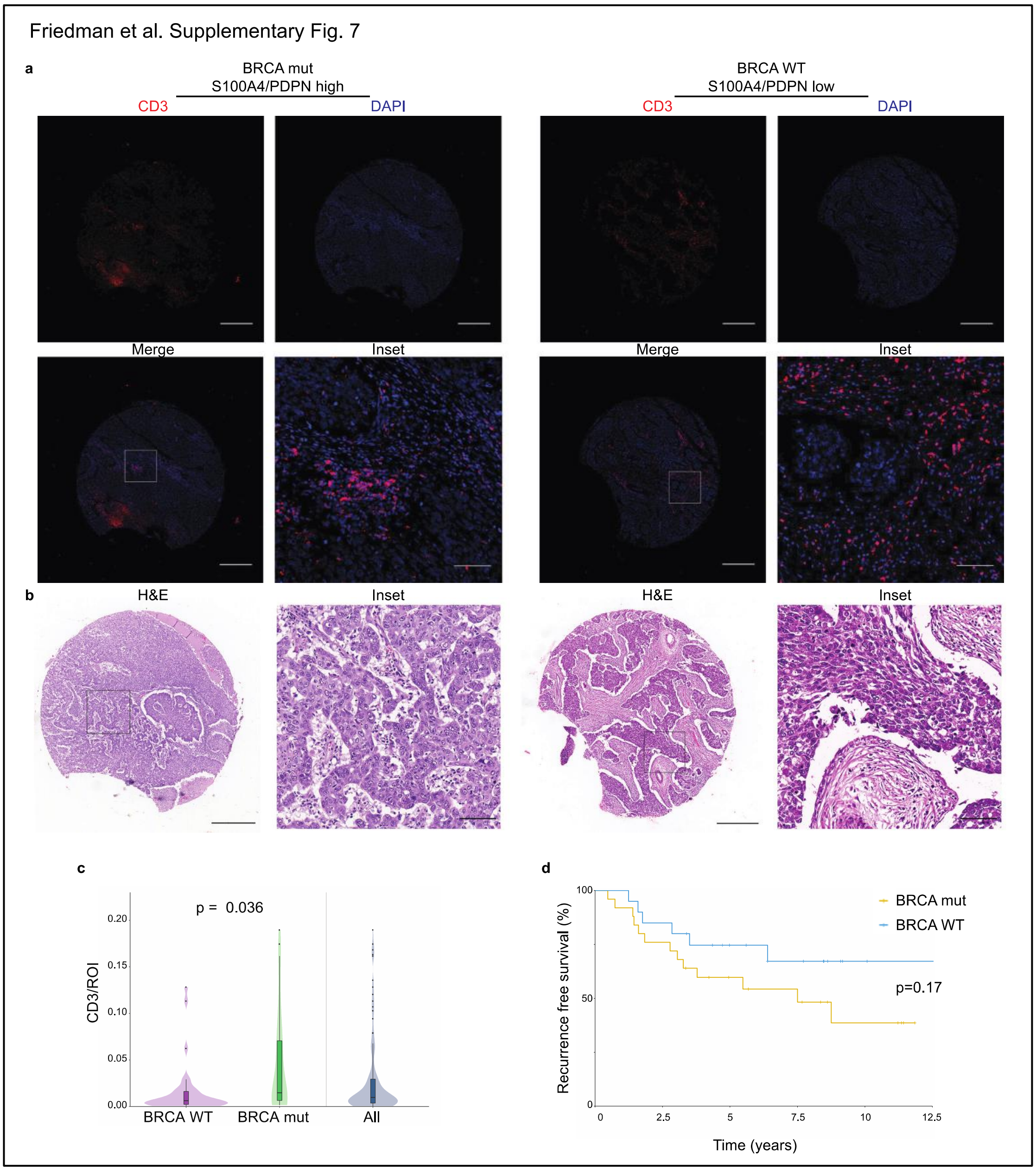


**Supplementary Fig. 7: BRCA status is not significantly correlated with recurrence free survival in a cohort of TNBC patients. a-b,** Representative images of CD3 and DAPI staining (**a**) and H&E stains (**b**) in a BRCA mutated (mut) patient and a BRCA WT patient from our cohort of 72 TNBC patients (Serial sections of the same cores used in Fig. 6a are shown). Scale bar = 500µm; inset scale bar = 80µm. **c,** Vase-box plot depicting CD3 staining scores (see Methods) in BRCA WT (n=25) *vs* BRCA mut (n=20) patients, as well as the total TNBC cohort (n=72). p Value was calculated using a student's t-test. **d,** 45 TNBC patients were stratified by BRCA mutational status and the association with recurrence free survival was assessed by KM analysis. P-value was calculated using log rank test.

**Supplementary Table 1.** Summary of biological samples

| Mouse ID | Tissue | Batch num | n cells |
|---|---|---|---|
| m1 | Normal mammary fat pad | 1 | 207 |
| m1 | Normal mammary fat pad | 2 | 143 |
| m1 | Normal mammary fat pad | 3 | 209 |
| m2 | Normal mammary fat pad | 4 | 204 |
| m2 | Normal mammary fat pad | 5 | 174 |
| m2 | Normal mammary fat pad | 6 | 114 |
| m3 | Normal mammary fat pad | 7 | 188 |
| m3 | Normal mammary fat pad | 8 | 92 |
| m3 | Normal mammary fat pad | 9 | 195 |
| m4 | Primary tumor, 2 weeks post injection | 10 | 243 |
| m4 | Primary tumor, 2 weeks post injection | 11 | 233 |
| m5 | Primary tumor, 2 weeks post injection | 12 | 255 |
| m5 | Primary tumor, 2 weeks post injection | 13 | 244 |
| m5 | Primary tumor, 2 weeks post injection | 14 | 243 |
| m6 | Primary tumor, 2 weeks post injection | 15 | 213 |
| m6 | Primary tumor, 2 weeks post injection | 16 | 167 |
| m7 | Primary tumor, 4 weeks post injection | 17 | 96 |
| m7 | Primary tumor, 4 weeks post injection | 18 | 60 |
| m7 | Primary tumor, 4 weeks post injection | 19 | 65 |
| m7 | Primary tumor, 4 weeks post injection | 20 | 38 |
| m7 | Primary tumor, 4 weeks post injection | 21 | 60 |
| m8 | Primary tumor, 4 weeks post injection | 22 | 218 |
| m8 | Primary tumor, 4 weeks post injection | 23 | 193 |
| m8 | Primary tumor, 4 weeks post injection | 24 | 231 |
| m8 | Primary tumor, 4 weeks post injection | 25 | 229 |
| m8 | Primary tumor, 4 weeks post injection | 26 | 236 |
| m8 | Primary tumor, 4 weeks post injection | 27 | 237 |
| m9 | Primary tumor, 4 weeks post injection | 28 | 144 |
| m9 | Primary tumor, 4 weeks post injection | 29 | 144 |
| m9 | Primary tumor, 4 weeks post injection | 30 | 116 |
| m9 | Primary tumor, 4 weeks post injection | 31 | 129 |
| m10 | Primary tumor, 4 weeks post injection | 32 | 165 |
| m10 | Primary tumor, 4 weeks post injection | 33 | 136 |
| m11 | Primary tumor, 4 weeks post injection | 34 | 81 |
| m11 | Primary tumor, 4 weeks post injection | 35 | 156 |
| m11 | Primary tumor, 4 weeks post injection | 36 | 39 |
| m11 | Primary tumor, 4 weeks post injection | 37 | 136 |
| m12 | Primary tumor, 4 weeks post injection | 38 | 92 |
| m12 | Primary tumor, 4 weeks post injection | 39 | 121 |
| m12 | Primary tumor, 4 weeks post injection | 40 | 219 |
| m12 | Primary tumor, 4 weeks post injection | 41 | 258 |
| m13 | Lung metastases | 42 | 128 |
| m13 | Lung metastases | 43 | 112 |
| m13 | Lung metastases | 44 | 118 |
| m14 | Lung metastases | 45 | 217 |
| m14 | Lung metastases | 46 | 198 |
| m14 | Lung metastases | 47 | 212 |
| m15 | Lung metastases | 48 | 200 |
| m15 | Lung metastases | 49 | 92 |
| m15 | Lung metastases | 50 | 33 |

**Supplementary Table 5.** Top 40 DE genes between EpCAM-GFP- sCAFs and EpCAM+GFP+ 4T1 cancer cells.

| * Gene | **GFP-EpCAM- | **GFP-EpCAM+ | **GFP+EpCAM- | **GFP+EpCAM+ |
|---|---|---|---|---|
| Ccl3 | 12.4057137 | 7.53736747 | 0 | 0 |
| Hdc | 11.4576988 | 5.92221425 | 2.49605844 | 0 |
| Ccl4 | 13.1592977 | 8.0952709 | 0.94731761 | 1.74952899 |
| Serpina1b | 10.4637127 | 2.77549865 | 1.51428295 | 0 |
| Il13 | 9.99739237 | 3.66610358 | 0 | 0 |
| Osm | 9.88473918 | 3.37231169 | 0.94731761 | 0 |
| Ramp2 | 11.1172711 | 5.21925309 | 3.36216697 | 1.36456784 |
| Cyp11a1 | 9.74972947 | 4.11868813 | 0 | 0 |
| Des | 10.5811615 | 3.66610358 | 1.92023697 | 0.83793993 |
| Ifitm1 | 10.4542694 | 3.66610358 | 2.2366874 | 0.83793993 |
| Csf2rb | 10.4174865 | 4.53723893 | 2.2366874 | 0.83793993 |
| Cyp4b1 | 9.51858347 | 2.77549865 | 0.94731761 | 0 |
| Gpx3 | 11.5223889 | 11.0727436 | 1.92023697 | 2.05310563 |
| Serping1 | 11.5042001 | 4.67569914 | 3.36216697 | 2.05310563 |
| Csf2rb2 | 10.7807219 | 4.67569914 | 1.51428295 | 1.36456784 |
| Col3a1 | 11.0819377 | 4.67569914 | 2.49605844 | 1.74952899 |
| Cdh13 | 9.2649728 | 5.02567036 | 2.2366874 | 0 |
| Meg3 | 9.24685513 | 1.73935152 | 0 | 0 |
| Cxcl12 | 9.21738513 | 4.46268418 | 0 | 0 |
| Il4 | 9.16431816 | 3.79323816 | 0 | 0 |
| Etv4 | 6.41711128 | 8.55157789 | 9.2808219 | 9.43222119 |
| Krt14 | 7.75474804 | 10.9561468 | 7.87607232 | 10.7783469 |
| Rhox5 | 6.39095167 | 8.97014622 | 7.20622205 | 9.44694654 |
| C3 | 7.71312431 | 9.88553719 | 9.87716676 | 10.7757571 |
| Msln | 6.41711128 | 8.88349371 | 9.30642962 | 9.48232149 |
| Mmp13 | 6.30950041 | 7.95952768 | 8.56179708 | 9.40231178 |
| Tslp | 6.22317397 | 9.24776549 | 8.36462742 | 9.34056589 |
| Padi4 | 6.41711128 | 9.44461747 | 9.71453541 | 9.64157013 |
| Il24 | 5.96399668 | 8.49658613 | 8.3275886 | 9.19308659 |
| Lad1 | 6.7199833 | 9.3978649 | 9.15726209 | 9.95142293 |
| Sfn | 6.5172496 | 10.8218544 | 7.23317612 | 9.74977814 |
| Cldn4 | 8.13171734 | 11.5909493 | 8.61400156 | 11.3842299 |
| Kcnn4 | 6.5648274 | 9.5085525 | 8.96698651 | 9.82810914 |
| Ankrd1 | 6.0993976 | 8.76051157 | 9.22596642 | 9.3905108 |
| Mmp9 | 8.52956968 | 11.4795616 | 10.8115591 | 11.8584689 |
| Spp1 | 10.23835 | 12.8797319 | 13.7380379 | 13.6181744 |
| Tns4 | 6.33716494 | 8.93265869 | 9.96152033 | 9.79011695 |
| Anxa8 | 6.03328503 | 9.38530556 | 8.7320854 | 9.49339766 |
| Syt8 | 6.06671999 | 8.89771264 | 9.67248442 | 9.55057096 |
| Gpa33 | 6.84086377 | 9.75884732 | 9.66425526 | 10.3350618 |

* 20 most upregulated and 20 most downregulated genes from all genes with a minimum of 2^8 average normalized reads (between GFP+EpCAM+ and GFP-EpCAM-; 1525 in total). Genes upregulated in EpCAM-GFP- are in orange; Genes upregulated in EpCAM+GFP+ are in blue.

** Reads are presented as log2 normalized values.

**Supplementary Table 7**. TNBC cohort clinical characteristics

| Characteristic | | |
|---|---|---|
| n | | 72 |
| Age at Dx (mean) | | 55.97 |
| Tumor size, n (%)* | T1 | 41 (60.3) |
| | T2 | 23 (33.8) |
| | T3 | 4 (5.9) |
| LN, n (%) | Negative | 50 (72.5) |
| | Positive | 19 (27.5) |
| Grade, n (%) | 1 | 1 (1.4) |
| | 2 | 9 (12.9) |
| | 3 | 60 (85.7) |
| BRCA, n (%) | WT | 25 (34.7) |
| | Mut | 20 (27.8) |
| | Unknown | 27 (37.5) |

**Supplementary Table 8**. TNBC cohort univariate analyses

| Variable (n) | | Hazard Ratio | 95% CI | P-Value |
|---|---|---|---|---|
| *CK (70) | | 36514.700 | | 0.028 |
| *Pdpn (70) | | 14019.614 | | 0.013 |
| S100A4 (70) | | 0.058 | 0 - 10075.709 | 0.644 |
| CD3 (68) | | 4.539 | 0.058 - 356.571 | 0.497 |
| S100A4/Pdpn (70) | | 0.388 | 0.199 - 0.758 | 0.005 |
| Pdpn/CK (70) | | 0.956 | 0.861 - 1.065 | 0.424 |
| S100A4/CK (70) | | 0.909 | 0.791 - 1.045 | 0.180 |
| LN (69) | | 1.865 | 0.865 - 3.964 | 0.105 |
| **Tumor size (69) | T1 | 1 [Reference] | | |
| | T2 | 2.100 | 0.956 - 4.607 | 0.065 |
| | T3 | 4.475 | 1.258 - 15.921 | 0.021 |
| BRCA status (45) | WT | 1 [Reference] | | |
| | Mut | 0.624 | 0.315 - 1.238 | 0.177 |
| Age (72) | | 1.022 | 0.999 - 1.045 | 0.062 |

**Supplementary Table 10.** TNBC cohort Multiplicative Multivariate Analysis

| Variable (n) | Hazard Ratio | 95% CI | P-Value | Likelihood ratio test |
|---|---|---|---|---|
| S100A4/Pdpn (45) | 0.828 | 0.421 - 1.631 | 0.585 | |
| BRCA status (45) | 37.101 | 4.000 - 344.491 | 0.001 | |
| S100A4/Pdpn:BRCA (45) | 0.015 | 0.001 - 0.230 | 0.002 | < 0.001 |

| Variable (n) | Hazard Ratio | 95% CI | P-Value | Likelihood ratio test |
|---|---|---|---|---|
| Pdpn/Total ROI (45) | 5.57E+05 | 0.6 - 5.058E+11 | 0.059 | |
| BRCA status (45) | 0.143 | 0.021 - 0.9692 | 0.046 | |
| Pdpn/Total ROI:BRCA (45) | 8.26E+12 | 2.778 - 2.45E+18 | 0.039 | 0.095 |

**Supplementary Table 11.** Antibodies and reagents

| Antibody | Conjugation | Clone | Source | Cat# | Target species | Application |
|---|---|---|---|---|---|---|
| EpCAM | Alexa flour 488 | G8.8 | Biolegend | 118210 | mouse | FACS analysis and sorting |
| EpCAM | FITC | G8.8 | eBioscience | 11579182 | mouse | FACS analysis and sorting |
| CD45 | BV711 | 30-F11 | Biolegend | 103147 | mouse | FACS analysis and sorting |
| CD45 | APC | 30-F11 | Biolegend | 103112 | mouse | FACS analysis and sorting |
| Ter119 | Pacific blue | Ter119 | Biolegend | 116232 | mouse | FACS analysis and sorting |
| PDPN | APC | 8.1.1 | Biolegend | 127410 | mouse | FACS analysis and sorting |
| PDPN | APC/Cy7 | 8.1.1 | Biolegend | 127417 | mouse | FACS analysis and sorting |
| PDPN | biotin | 8.1.1 | Biolegend | 127404 | mouse | FACS analysis and sorting |
| Propidium iodide | - | - | Sigma Aldrich | P4170 | mouse | FACS analysis and sorting |
| S100A4 | - | polyclonal | Abcam | ab41532 | mouse, human | Immunofluoresence |
| PDPN | - | polyclonal | R&D Systems | AF3244 | mouse | Immunofluoresence |
| PDPN | - | D2-40 | Biolegend | 916605 | human | Immunofluoresence |
| Cytokeratin 7 | - | EPR17078 | Abcam | ab181598 | mouse | Immunofluoresence |
| Pan-cytokeratin | - | AE1/AE3 | Dako | M3515 | human | Immunofluoresence |
| DAPI | - | - | Biolegend | 422801 | - | Immunofluoresence |
| CD3 | - | D4V8L | Cell Signaling | 99940 | human | Immunofluoresence |
| CD45 | - | D9M8I | Cell Signaling | 13917 | human | Immunofluoresence |
| Ly-6C | PerCP/Cy5.5 | HK 1.4 | Biolegend | 128011 | mouse | FACS analysis and sorting |
| I-A/I-E | APC/Cy7 | M5/114.15.2 | Biolegend | 107628 | mouse | FACS analysis and sorting |
| HLA-DR | - | EPR3692 | Abcam | ab92511 | human | Immunofluoresence |
| $\alpha$-SMA | - | 1A4 | Sigma Aldrich | A2547 | huma/mouse | Immunofluoresence |
| $\alpha$-SMA | FITC | 1A4 | Sigma Aldrich | F3777 | huma/mouse | FACS analysis |
| NT5E/CD73 | - | D7F9A | Cell Signaling | 13160 | human | Immunofluoresence |
| Bovine anti-Goat | HRP | polyclonal | Jackson | 805-035-180 | goat | Immunofluoresence |
| Goat anti-Rabbit | HRP | polyclonal | Jackson | 111-035-144 | rabbit | Immunofluoresence |
| Goat anti-Mouse | HRP | polyclonal | Abcam | 97040 | mouse | Immunofluoresence |
| Opal 520 Reagent | Opal 520 | - | PE | FP1487001KT | - | Immunofluoresence |
| Opal 570 Reagent | Opal 570 | - | PE | FP1488A | - | Immunofluoresence |
| Opal 620 Reagent | Opal 620 | - | PE | FP1495 | - | Immunofluoresence |
| Opal 650 Reagent | Opal 650 | - | PE | FP1496A | | Immunofluoresence |
| Opal 690 Reagent | Opal 690 | | PE | FP1497A | - | Immunofluoresence |
| 1X Plus Amp Diluent | - | - | PE | FP1498 | - | Immunofluoresence |

**Supplementary Table 12**. Selected genes, priority, and fold change threshold for subset specific gene expression analysis

| Group | Gene | Priority | T_fold |
|---|---|---|---|
| Wound healing | Acta2 | 6 | 4.1 |
| Immune regulatory E | Cxcl12 | 9 | 4.9 |
| ECM | Mfap5 | 5 | 8.2 |
| NMF | Gsn | 5 | 18 |
| Inflammatory A | Cxcl1 | 4 | 3.5 |
| Immune regulatory L | Saa3 | 7 | 15 |
| Inflammatory B | Il6 | 17 | 5 |
| Antigen presentation | Spp1 | 7 | 3.1 |
| Protein folding | S100a4 | 13 | 1.77 |
